# Inter-trial effects in visual pop-out search: Factorial comparison of Bayesian updating models

**DOI:** 10.1101/353003

**Authors:** Fredrik Allenmark, Hermann J. Müller, Zhuanghua Shi

**Author notes:** Correspondence concerning this article should be addressed to Zhuanghua Shi, General and Experimental Psychology, Department of Psychology, Ludwig-Maximilians-Universität München, 80802, Munich, Germany.

## Abstract

Many previous studies on visual search have reported inter-trial effects, that is, observers respond faster when some target property, such as a defining feature or dimension, or the response associated with the target repeats versus changes across consecutive trial episodes. However, what processes drive these inter-trial effects is still controversial. Here, we investigated this question using a combination of Bayesian modeling of belief updating and evidence accumulation modeling in perceptual decision-making. In three visual singleton (‘pop-out’) search experiments, we explored how the probability of the response-critical states of the search display (e.g., target presence/absence) and the repetition/switch of the target-defining dimension (color/ orientation) affect reaction time distributions. The results replicated the mean reaction time (RT) inter-trial and dimension repetition/switch effects that have been reported in previous studies. Going beyond this, to uncover the underlying mechanisms, we used the Drift-Diffusion Model (DDM) and the Linear Approach to Threshold with Ergodic Rate (LATER) model to explain the RT distributions in terms of decision bias (starting point) and information processing speed (evidence accumulation rate). We further investigated how these different aspects of the decision-making process are affected by different properties of stimulus history, giving rise to dissociable inter-trial effects. We approached this question by (i) combining each perceptual decision making model (DDM or LATER) with different updating models, each specifying a plausible rule for updating of either the starting point or the rate, based on stimulus history, and (ii) comparing every possible combination of trial-wise updating mechanism and perceptual decision model in a factorial model comparison. Consistently across experiments, we found that the (recent) history of the response-critical property influences the initial decision bias, while repetition/switch of the target-defining dimension affects the accumulation rate, likely reflecting an implicit ‘top-down’ modulation process. This provides strong evidence of a disassociation between response- and dimension-based inter-trial effects.

## Introduction

In everyday life, we are continuously engaged in selecting visual information to achieve our action goals, as the amount of information we receive at any time exceeds the available processing capacity. The mechanisms mediating attentional selection enable us to act efficiently by prioritizing task-relevant, and deprioritizing irrelevant, information. Of importance for the question at issue in the present study, the settings that ensure effective action in particular task episodes are, by default, buffered by the attentional control system and carried over to subsequent task episodes, facilitating performance if the settings are still applicable and, respectively, impairing performance if they no longer apply owing to changes in the task situation (in which case the settings need to be adapted accordingly). In fact, in visual search tasks, such automatic carry-over effects may account for more of the variance in the response times (RTs) than deliberate, top-down task set [1]. A prime piece of evidence in this context is visual search for so-called singleton targets, that is, targets defined by being unique relative to the background of non-target (or distractor) items, whether they differ from the background by one unique feature (simple feature singletons) or a unique conjunction of features (conjuction singletons): singleton search is expedited (or slowed) when critical properties of the stimuli repeat (or change) across trials. Such inter-trial effects have been found for repetitions/switches of, for example, the target-defining color [2,3], size [4], position [5], and, more generally, the target-defining feature dimension [6,7]. The latter has been referred to as the dimension repetition/switch effect, that is: responding to a target repeated from the same dimension (e.g., color) is expedited even when the precise target feature is different across trials (e.g., changing from blue on one trial to red on the next), whereas a target switch from one dimension to another (e.g., from orientation to color) causes a reaction time cost (‘dimension repetition effect’, DRE) [8–10].

While inter-trial effects have been extensively studied, the precise nature of the processes that are being affected remains unclear. Much of the recent work has been concerned with the issue of the processing stage(s) at which inter-trial effects arise (for a review, see [11]). Müller and colleagues proposed that inter-trial effects, in particular the dimension repetition effect, reflect facilitation of search processes prior to focal-attentional selection (at a pre-attentive stage of saliency computation) [10]. However, using a non-search paradigm with a single item presented at a fixed (central) screen location, Mortier et al. [12] obtained a similar pattern of inter-trial effects – leading them to conclude that the DRE arises at the post-selective stage of response selection. Rangelov and colleagues [13] demonstrated that DRE effects can originate from distinct mechanisms in search tasks making different task demands (singleton feature detection and feature discrimination): pre-attentive weighting of the dimension-specific feature contrast signals and post-selective stimulus processing – leading them to argue in favor of a ‘multiple weighting systems hypothesis’. Based on the ‘priming of pop-out’ search paradigm, a similar conclusion [11] has also been proposed, namely, inter-trial effects arise from both attentional selection and post-selective retrieval of memory traces from previous trials [4,14], favoring a dual-stage account [15].

It is important to note that those studies adopted very different paradigms and tasks to examine the origins of inter-trial effects, and their analyses are near-exclusively based on differences in mean RTs. Although such analyses are perfectly valid, much information about trial-by-trial changes is lost. Recent studies have shown that the RT distribution imposes important constraints on theories of visual search [16,17]. RT distributions in many different task domains have been successfully modeled as resulting from a process of evidence accumulation [18,19]. One influential evidence accumulation model is the drift-diffusion model (DDM) [20–22]. In the DDM, observers sequentially accumulate multiple pieces of evidence, each in the form of a log likelihood ratio of two alternative decision outcomes (e.g., target present vs. absent), and make a response when the decision information reaches a threshold (see Figure 1). The decision process is governed by three distinct components: a tendency to drift towards either boundary (drift rate), the separation between the decision boundaries (boundary separation), and a starting point. These components can be estimated for any given experimental condition and observer by fitting the model to the RT distribution obtained for that condition and observer.

Estimating these components makes it possible to address a question that is related to, yet separate from the issue of the critical processing stage(s) and that has received relatively less attention: do the faster RTs after stimulus repetition reflect more efficient stimulus processing, for example: expedited guidance of attention to more informative parts of the stimulus, or rather a bias towards giving a particular one of the two alternative responses or, respectively, a tendency to require less evidence before issuing either response. The first possibility, more efficient processing, would predict an increase in the drift rate, that is, a higher speed of evidence accumulation. A bias towards one response or a tendency to require less evidence would, on the other hand, predict a decreased distance between the starting point and the decision boundary associated with that response. In the case of bias, this would involve a shift of the starting point towards that boundary, while a tendency to require less evidence would be reflected in a decrease of the boundary separation. While response bias is more likely associated with changes at the post-selective (rather than pre-attentive) processing stage, the independence of the response selection and the attentional selection stage has been challenged [23].

For simple motor latencies and simple-detection and pop-out search tasks [24], there is another parsimonious yet powerful model, namely the LATER (Linear Approach to Threshold with Ergodic Rate) model [25,26]. Unlike the drift-diffusion model, which assumes that evidence strength varies across the accumulative process, the LATER model assumes that evidence is accumulated at a constant rate during any individual perceptual decision, but that this rate varies randomly across trials following a normal distribution (see Figure 1). Such a pattern has been observed, for instance, in the rate of build-up of neural activity in the motor cortex of monkeys performing a saccade-to-target task [27]. Similar to the DDM, the LATER model has three important parameters: the ergodic rate (*r*), the boundary separation (*θ*), and a starting point (*S*_0_). However, the boundary separation and starting point are not independent, since the output of the model is completely determined by the rate and the separation between the starting point and the boundary; thus, in effect, the LATER model has only two parameters.

The evidence accumulation process can be interpreted in terms of Bayesian probability theory [26,28]. On this interpretation, the ‘linear approach to threshold with ergodic rate’ represents the build-up of the posterior probability that results from adding up the log likelihood ratio (i.e., ‘evidence’) of a certain choice being the correct one and the initial bias that derives from the prior probability of two choices. The prior probability should affect the starting point *S*_0_ of the evidence accumulation process: *S*_0_ should be the closer to the boundary the higher the prior probability of the outcome that boundary represents. The drift rate, by contrast, should be influenced by any factor that facilitates or impedes efficient accumulation of task-relevant sensory evidence, such as spatial attentional selection.

**Figure 1.**
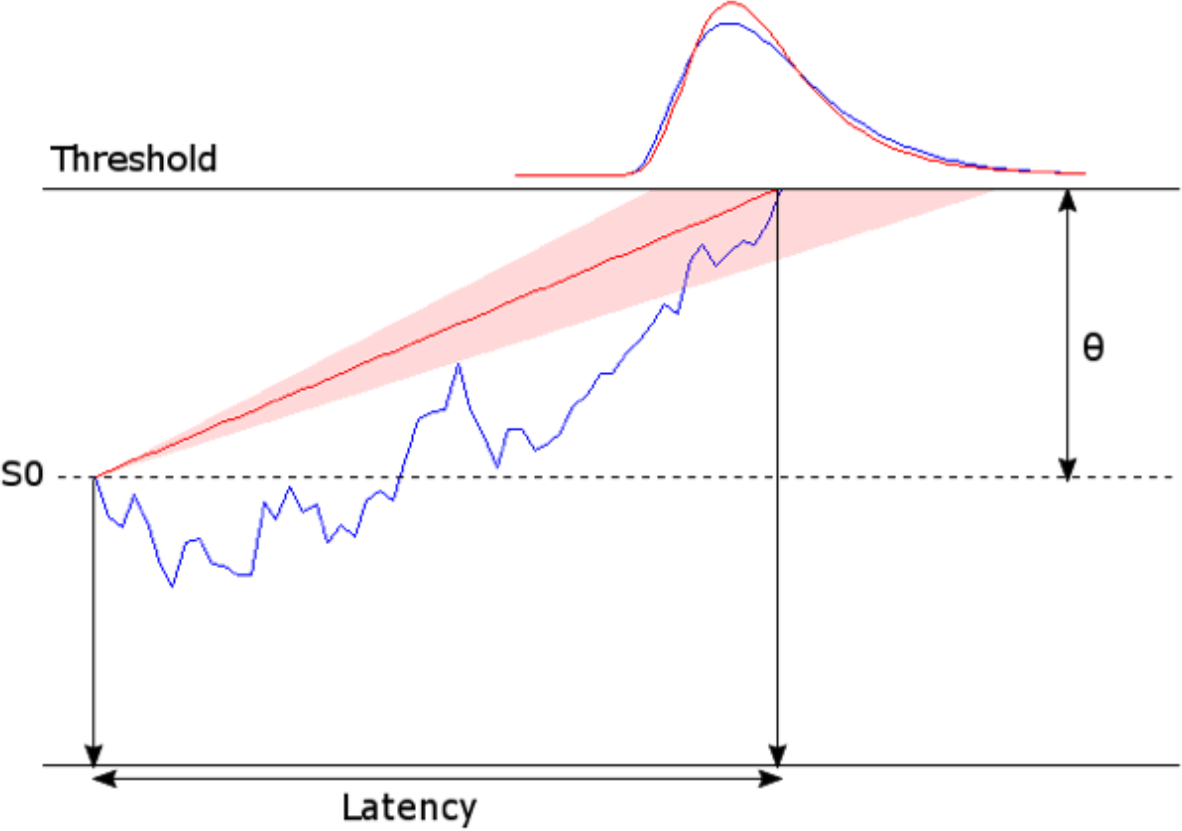
Illustrations of the drift diffusion model (DDM, shown in blue) and the LATER model (shown in red). The DDM assumes that evidence accumulates, from the starting point (S_0_),through random diffusion in combination with a drift rate r until a boundary (i.e., threshold, θ) is reached. The LATER model makes the same assumptions, except that the rater is considered to be constant within any individual trial, but to vary across trials (so as to explain trial-to-trial variability in RTs).

The present study was designed to clarify the nature of the inter-trial effects for manipulations of target presence and the target-defining dimension as well as inter-trial dimension repetitions and switches. If inter-trial effects reflect a decision bias, this should be reflected in changes of the decision boundary and/or the starting point. By contrast, if inter-trial effects reflect changes in processing efficiency, which might result from allocating more attentional resources (or ‘weight’) to the processing of the repeated feature/dimension [6], the accumulation rate *r* should be changed. Note that neither the DDM nor the LATER model provides any indication of how the initial starting point might change across trials. Given that the inter-trial effects are indicative of the underlying trial-by-trial dynamics, we aimed to further analyze trial-wise changes of the prior and the accumulation rate, and examine how a new prior is learned when the stimulus statistics change, as reflected in changes of the starting point to decision boundary separation during the learning process.

To address these inter-trial dynamics, we adopted the Dynamic Belief Model (DBM) [29]. The DBM has been successfully used to explain why performance on many tasks is better when a stimulus matches local patterns in the stimulus history even in a randomized design where it is not actually possible to use stimulus history for (better-than-chance) prediction. Inter-trial effects arise naturally in the DBM. This is because the DBM assumes a prior belief about non-stationarity, that is: participants are updating their beliefs about the current stimulus statistics while assuming that these can change at any time. The assumption of non-stationarity leads to something similar to exponential discounting of previous evidence, that is, the weight assigned to previous evidence decreases exponentially with the time (or number of updating events) since it was acquired. Consequently, current beliefs about what is most likely to happen on an upcoming trial will always be significantly influenced by what occurred on the previous trial, resulting in intertrial effects. Thus, here we combine a belief-updating model closely based on the DBM, for modelling the learning of the prior, with the DDM and, respectively, the LATER model for predicting RTs. A very similar model has previously been proposed to explain results in saccade-to-target experiments [30]. We also consider the possibility that the evidence accumulation rate as well as the starting point may change from trial to trial.

To distinguish between different possible ways in which stimulus history could have an influence via updating of the starting point and/or the rate, we performed three visual search experiments, using both a detection and a discrimination task and manipulating the probability of target presence, as well as the target-defining dimension. Based on the RT data, we then performed a factorial model comparison (cf. [31]), where both the response history and the history of the target dimension can affect either the starting point or the rate. The results show that the model that best explains both the effects of our probability manipulation and the inter-trial effects is the one in which the starting point is updated based on response history and the rate is updated based on the history of the target dimension.

## Results

Experiments 1 and 2 both consisted of three equally long blocks. The frequency of pop-out target presence (or absence) was varied across blocks in Experiment 1. In Experiment 2, a target was always present, and the frequency of the target being a color-defined or, alternatively, an orientation-defined singleton was varied across blocks. In Experiment 3, target presence and absence were kept equally frequent, as were trials with color- and orientation-defined singleton targets. One implication of this design is that the high-frequency condition for one target condition (present/absent, color/orientation) was implemented in the same block as the low-frequency condition for the other target condition. So, in all figures and analyses of the effects of frequency, the high- and low-frequency conditions are based on data collected in different blocks for each target condition, while the data for the medium-frequency condition comes from the same block for each target condition.

### Error rates

**Figure 2.**
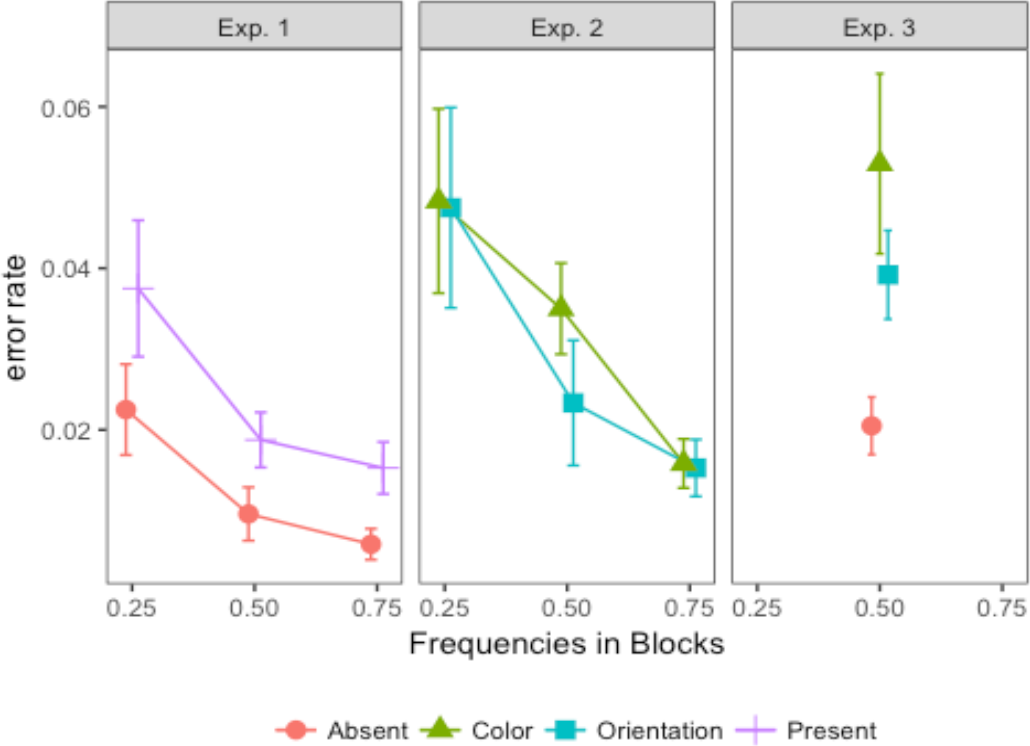
Error rates in Experiments 1, 2, and 3, for all combinations of target frequency. Target frequency is defined relative to the target condition, as the frequency with which that target condition occurred within a given block. This means that, for a given frequency, the data from the different target conditions do not necessarily come from the same block of the experiment. Error bars show the standard error of the mean.

The singleton search was quite easy, with participants making few errors overall: mean error rates were 1.5%, 2.5%, and 3.3% in Experiments 1, 2, and 3 respectively (Figure 2). Despite the low average error rates, error rates differed significantly between blocks in both Experiments 1 and 2 [*F*(1.34,14.78) = 11.50, *p* < 0.01,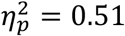, *BF* = 8372, and *F*(2,22) = 12.20, *p* < 0.001,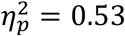, *BF* = 3729, respectively]: as indicated by post-hoc comparisons (Supplement S1), error rates were higher in the low-frequency blocks compared to the medium- and high-frequency blocks, without a significant difference between the latter. In addition, in Experiment 1, error rates were overall higher for target-present than for target-absent trials, that is, there were more misses than false alarms, *F*(1,11) = 11.43, *p* < 0.01,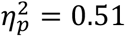 *BF* = 75. In contrast, there was no difference in error rates between color and orientation targets in Experiment 2, *F*(1,11) = 0.70, *p* = 0.42, *BF* = 0.33. In Experiment 3, there was no manipulation of target (or dimension) frequency, but like in Experiment 1, error rates were higher on target-present than on target-absent trials, *t*(11) = 4.25, *p* < 0.01, *BF* = 30.7; and similar to Experiment 2, there was no significant difference in error rates between color and orientation targets, *t*(11) = 1.51, *p* = 0.16, *BF* = 0.71.

### Mean Reaction times (RTs)

**Figure 3.**
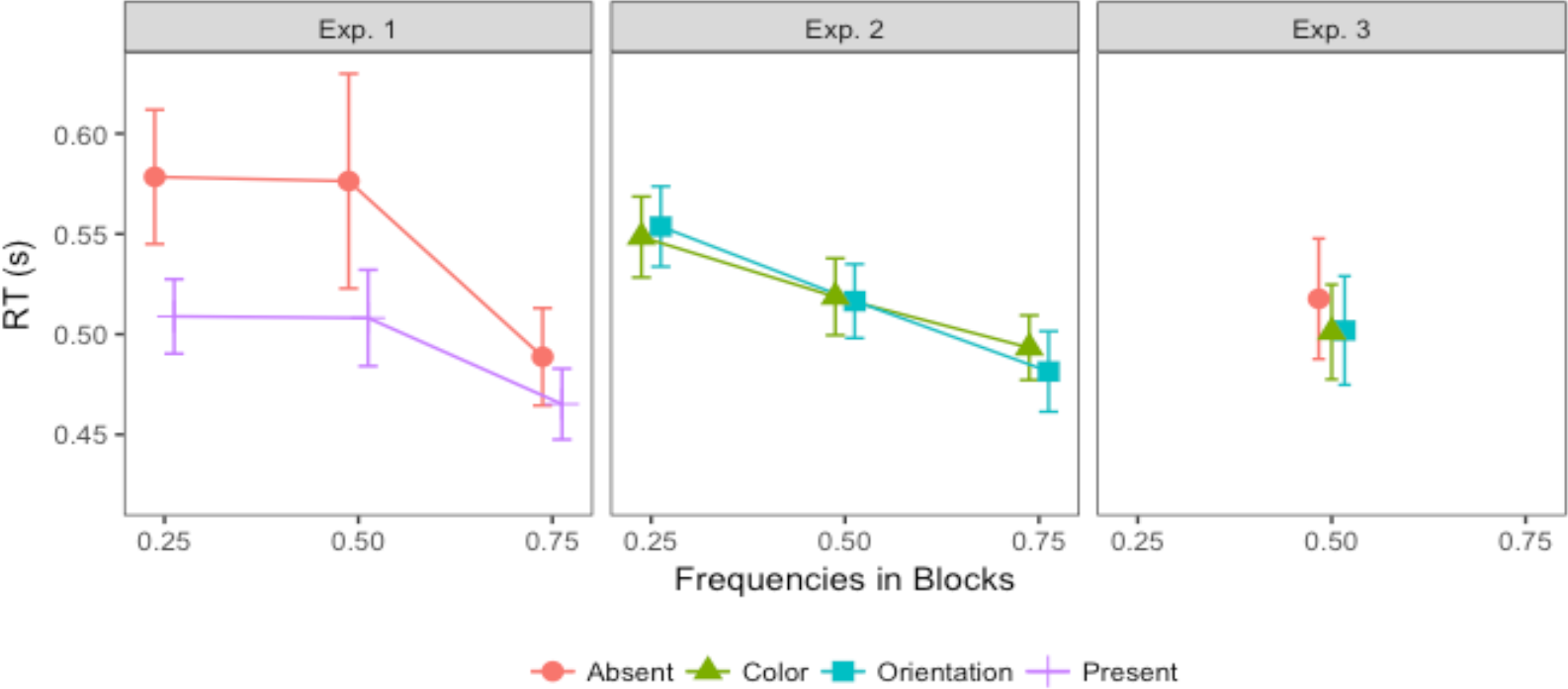
Mean RTs in Experiments 1, 2, and 3, for all combinations of target condition and target frequency. Target frequency is defined relative to the target condition, as the frequency with which that target condition occurred within a given block. This means that for a given frequency, the data from the different target conditions do not necessarily come from the same block of the experiment. Error bars show the standard error of the mean.

Given the low error rates, we analyzed only RTs from trials with a correct response, though excluding outliers, defined as trials on which the inverse RT (i.e., 1/RT) was more than three standard deviations from the mean for any individual participant. Figure 3 presents the pattern of mean RTs for all three experiments. In both Experiments 1 and 2, the main effect of frequency was significant [*F*(2,22) = 10.25, *p* < 0.001,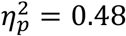, *BF* = 73, and, respectively, *F*(1.27,13.96) = 29.83, *p* < 0.01,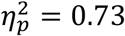, *BF* = 8.7 * 10^8^]. Post-hoc comparisons (see Supplement S2) confirmed RTs to be faster in high-frequency compared to low-frequency blocks, indicative of participants adapting to the stimulus statistics in a way such as to permit faster responses to the most frequent type of trial within a given block. In addition, in Experiment 1, RTs were faster for target-present than for target-absent trials [*F*(1,11) = 5.94, *p* < 0.05,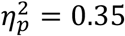, *BF* = 51], consistent with the visual search literature. In contrast, there was no difference between color- and orientation-defined target trials in Experiment 2, and no interaction between target condition and frequency in either Experiment 1 or 2 (Supplement S2) – suggesting that the effect of frequency is independent of the target stimuli.

Comparing the error rates depicted in Figure 2 and the mean RTs in Figure 3, error rates tended to be lower for those frequency conditions for which RTs were faster. While this rules out simple speed-accuracy trade-offs, it indicates that participants were adapting to the statistics of the stimuli in a way that permitted faster and more accurate responding to the most frequent type of trial within a given block, at the cost of slower and less accurate responding on the less frequent trial type. A possible explanation of these effects is a shift of the starting point of a drift-diffusion model towards the boundary associated with the response associated with the most frequent type of trial; as will be seen below (in the modeling section), the shapes of the RT distributions were consistent with this interpretation.

Without a manipulation of frequency, Experiment 3 yielded a standard outcome: all three types of trial yielded similar mean RTs, *F*(2,22) = 2.15, *p* = 0.14, *BF* = 0.71. This is different from Experiment 1, in which target-absent RTs were significantly slower than target-present RTs. This difference was likely obtained because the target-defining dimension was kept constant within short mini-blocks in Experiment 1, but varied randomly across trials in Experiment 3, yielding a dimension switch cost and therefore slower average RTs on target-present trials (see modeling section for further confirmation of this interpretation).

### Inter-trial effects

**Figure 4.**
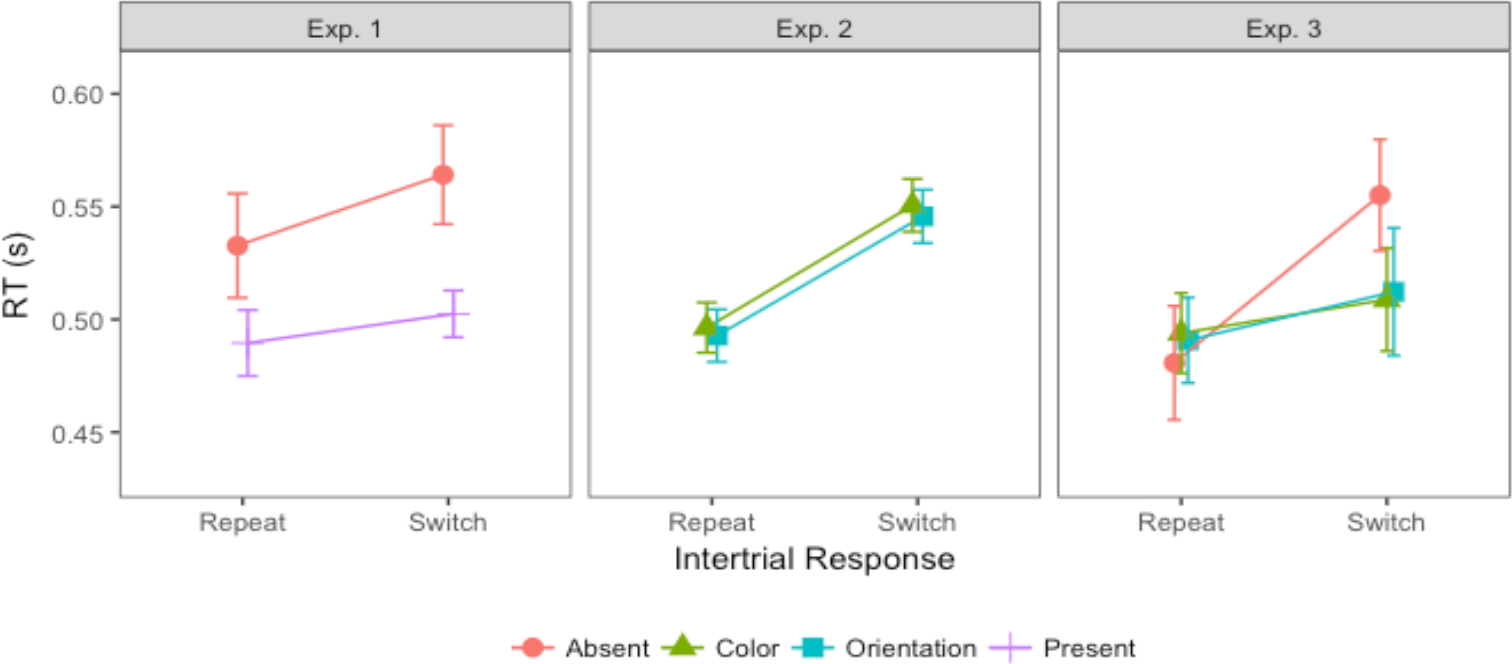
Inter-trial effects on mean RTs for all three experiments. Error bars show the standard error of the mean.

Given our focus on inter-trial dynamic changes in RTs, we compared trials on which the target condition was switched to trials on which it was repeated from the previous trial. Figure 4 illustrates the inter-trial effects for all three experiments. RTs were significantly faster on target-repeat than on target-switch trials, in all experiments: Experiment 1 [ *F*(1,11) = 6.13, *p* < 0.05,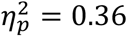, *BF* = 0.81], Experiment 2 [*F*(1,11) = 71.29, *p* < 0.001,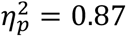, *BF* = 2.6 * 10^7^], and Experiment 3 [ *F*(1,11) = 32.68, *p* < 0.001,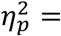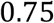, *BF* = 625]. Note that for Experiment 1, despite the significant target-repeat/switch effect, the ‘inclusion’ BF (see Methods) suggests that this factor is negligible compared to other factors; a further post-hoc comparison of repeat versus switch trials has a BF of 5.88, compatible with the ANOVA test. The target repetition effect in all three experiments is consistent with trial-wise updating of an internal model (see the modeling section). The target repetition/switch effect was larger for target-absent responses (i.e., comparing repetition of target absence to a switch from target presence to absence) than for target-present responses in Experiment 3 (interaction inter-trial condition x target condition, *F*(1,11) = 14.80, *p* < 0.01,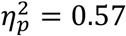, *BF* = 18), while there was no such a difference in Experiment 1, *F*(1,11) = 2.55, *p* = 0.14, *BF* = 0.43, and also no interaction between target dimension and inter-trial condition in Experiment 2, *F*(1,11) = 0.014, *p* = 0.91, *BF* = 0.76. These findings suggest that, while the target repetition/switch effect as such is stable across experiments, its magnitude may fluctuate depending on the experimental condition. The interaction between target condition and inter-trial condition seen in Experiment 3, but not in Experiment 1, is likely attributable to the fact that color and orientation targets were randomly interleaved in Experiment 3, so that target-present repetitions include trials on which the target dimension did either repeat or change – whereas the target dimension was invariably repeated on consecutive target-present trials in Experiment 1. The effects of repeating/switching the target dimension are considered further below.

Note that in all experiments, we mapped two alternative target conditions to two fixed alternative responses. The repetition and switch effects described above may be partly due to response repetitions and switches. To further examine dimension repetition/switch effects when both dimensions were mapped to the same response, we extracted those target-present trials from Experiment 3 on which a target was also present on the immediately preceding trial. Figure 5 depicts the mean RTs for the dimension-repeat versus-switch trials. RTs were faster when the target dimension repeated compared to when it switched, *F*(1,11) = 25.06, *p* < 0.001,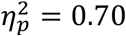, *BF* = 1905, where this effect was of a similar magnitude for color- and orientation-defined targets [interaction target dimension x dimension repetition, *F*(1,11) = 0.04, *p* = 0.84, *BF* = 0.33]. There was also no overall RT difference between the two types of target [main effect of target dimension, *F*(1,11) = 0.16, *p* = 0.69, *BF* = 0.34], indicating that the color and orientation targets were equally salient. This pattern of dimension repetition/switch effects is in line with the dimension-weighting account [8]. Of note, there was little evidence of a dimension repetition benefit from two trials back, that is, from trial *n*-2 to trial *n*: the effect was very small (3 ms) and not statistically significant [t(23)=0.81, p=0.43, BF=0.38].

**Figure 5.**
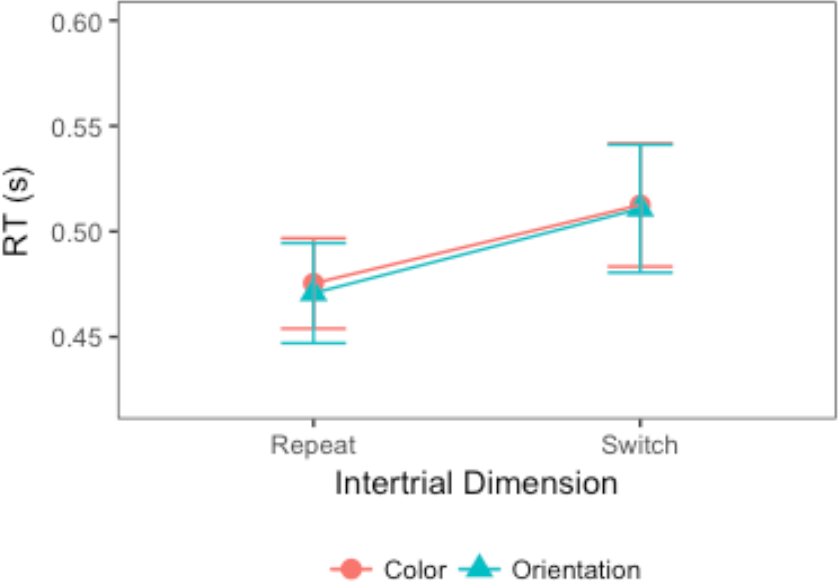
Dimension repetition/switch effect in Experiment 3. Mean RTs were significantly faster when the target-defining dimension was repeated. Error bars show the standard error of the mean.

In addition to inter-trial effects from repetition versus switching of the target dimension, there may also be effects of repeating/switching the individual target-defining features. To examine for such effects, we extracted those trials on which a target was present and the target dimension stayed the same as on the preceding trial, and examined them for (intra-dimension) target feature repetition/switch effects. See Figure 6 for the resulting mean RTs. In Experiments 1 and 3, there was no significant main effect of feature repetition/switch [Exp. 1:*F*(1,11) = 0.30, *p* = 0.593, *BF* = 0.30, Exp. 3: *F*(1,11) = 3.77, *p* = 0.078, *BF* = 0.76], nor was there an interaction with target dimension [Exp. 1: *F*(1,11) = 2.122, *p* = 0.17, BF = 0.44, Exp. 3: *F*(1,11) = 0.007, *p* = 0.93, *BF* = 0.38]. In contrast, in Experiment 2 (which required an explicit target dimension response), RTs were significantly faster when the target feature repeated compared to when it switched within the same dimension, *F*(1,11) = 35.535, *p* < 0.001,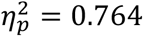, *BF* = 13, and this effect did not differ between the target-defining, color and orientation, dimensions, *F*(1,11) = 1.858, *p* = 0.2, *BF* = 0.57. Note though that, even in Experiment 2, this feature repetition/switch effect was smaller than the effect of dimension repetition/switch (20 vs. 54 ms, t(11)=5.20, p<0.001, BF=122).

**Figure 6.**
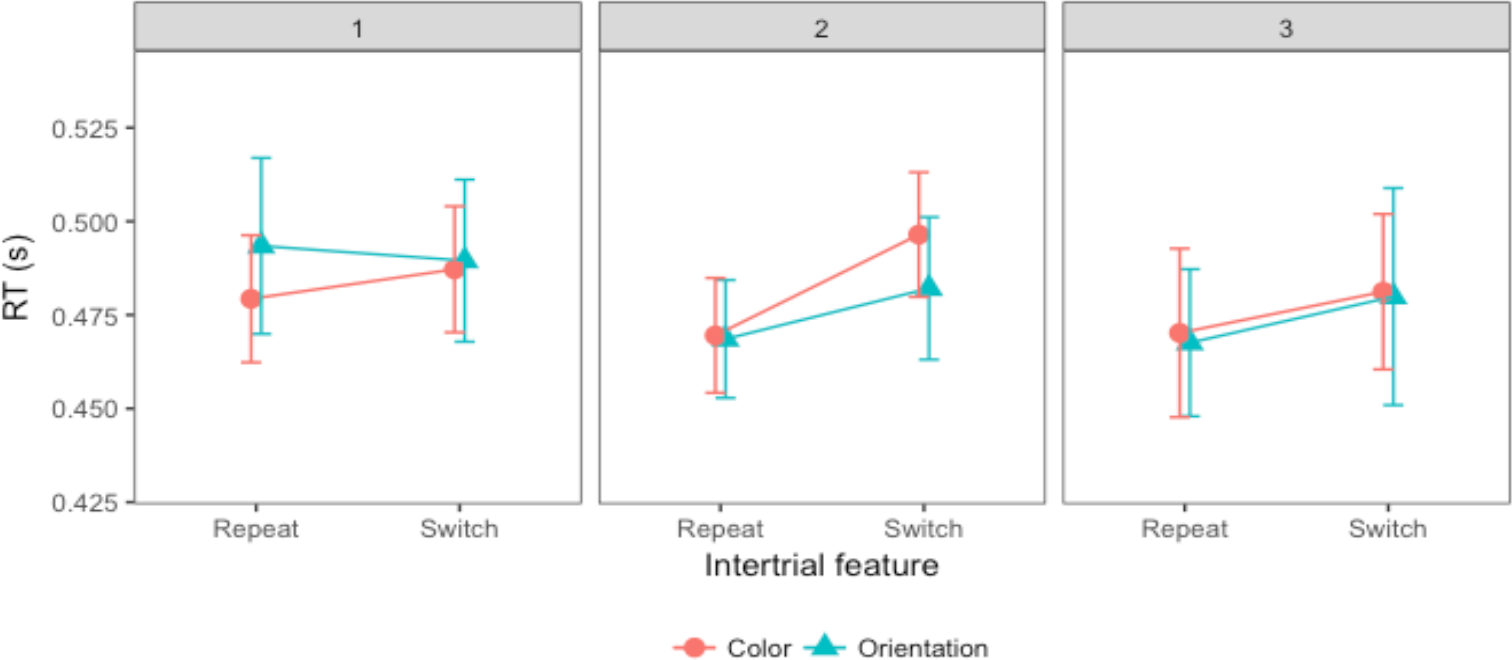
Feature repetition/switch effects on mean RTs for all three experiments. Error bars show the standard error of the mean.

In summary, the results revealed RTs to be expedited when target presence or absence or, respectively, the target-defining dimension (on target-present trials) was repeated on consecutive trials. However, the origin of these inter-trial effects is unclear: The faster RTs for cross-trial repetitions could reflect either more efficient stimulus processing (e.g., as a result of greater ‘attentional ‘weight’ being assigned to a repeated target dimension) or a response bias (e.g., an inclination to respond ‘target present’ based on less evidence on repeat trials), or both. In the next section, we will address the origin(s) of the inter-trial effects by comparing a range of generative computational models and determining which parameters are likely involved in producing these effects. Because feature-specific intertrial effects, if reliable at all (they were significant only in Exp. 2, which required an explicit target dimension response), were smaller than the inter-trial effects related to either target presence/absence or the target-defining dimension (e.g., in Exp. 3, a significant dimension-based inter-trial effect of 39 ms compares with a non-significant feature-based effect of 11 ms), we chose to ignore the feature-related effect in our modeling attempt.

### Dynamic Bayesian updating and inter-trial effects

#### Factorial comparison of multiple updating models

To identify the origins of the observed inter-trial effects, we systematically compared a multiplicity of computational models using the factorial comparison method [31]. Given that both the DDM and the LATER model provide a good prediction of the RT distributions, we consider the model of RT distributions as one factor (DDM vs. LATER).

Both models have the same parameters: the evidence accumulation rate (*r*), the initial starting point (*S*_0_), and the decision threshold (*θ*). The DDM model has one additional parameter: non-decision time (*T*_*er*_). Here we also added a non-decision time parameter to the LATER model, and considered the presence versus absence of a non-decision time as one factor (i.e., non-decision time fixed to zero vs. non-decision time as a free parameter).

**Table 1.** A list of the levels and the associated parameters for each of the four factors.

| FACTORS | LEVELS | PARAMETERS |
| --- | --- | --- |
| NON-DECISION TIME | Without | None |
| | With | $T_{er}$ |
| EVIDENCE ACCUMULATION MODEL | DDM | $r, T, \sigma$ |
| | LATER | $r, T, \sigma$ |
| RDF-BASED UPDATING<br>OR<br>TDD-BASED UPDATING | No updating | - |
| | $S_0$ full memory | $\beta_0$ |
| | $S_0$ with decay | $\alpha, \beta_0$ |
| | Binary rate | $\kappa$ |
| | Rate with decay | $\alpha, \Delta$ |
| | Weighted rate | $\alpha, \Delta$ |

One of the main purposes of the model comparison was to investigate through what mechanisms response history and the history of the target dimension influence RTs. To this end, we introduced the influence of the history of the ‘response-defining feature’ (RDF) and of the ‘target-defining dimension’ (TDD) on updating of the parameters of the RT distribution model as two separate factors. For each factor, we considered six different forms of updating (factor levels). **Error! Reference source not found.** lists all factor levels and the associated parameters for each of the four factors.

**Level 1 (No update)**: RDF/TDD repetition/switch does not affect any model parameters.

**Level 2** (*S*_0_ **with full memory)**: RDF/TDD repetition/switch updates the initial starting point (*S*_0_) according to the whole prior history. As suggested by [26] and [19], *S*_0_ is determined by the log prior odds of two decision outcomes (*H* vs. *~H*):

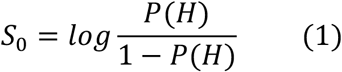

Here we assume that the prior probability *P*(*H*), rather than being fixed, is updated trial-wise according to Bayesian inference, because participants are learning the frequencies of different stimulus properties (such as target present/absent or color/orientation) and using this knowledge as a prior when making perceptual decisions. Thus, the posterior of the prior is:

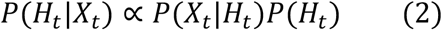

This updating can be modeled by using a Beta distribution as the starting distribution on the prior (a hyperprior) and updating after each trial using the Bernoulli likelihood. We assume that participants were unbiased at the beginning of the experiment (i.e., the two parameters of the Beta distribution initially had the same value *β*_0_) and gradually updated their prior based on the trial history. The updating fully determines the starting point on each trial based on the stimulus history and the shape of the starting distribution (determined by *β*_0_); accordingly, the shape parameter of the starting distribution, *β*_0_, is the only free parameter. Figure 7 illustrates the updating.

For updating based on the RDF, a single prior *p* is being learned, representing the probability of target-present trials (with the probability of a target-absent trial being 1 − *p*). For updating based on the history of the TDD, we assume a separate prior is being learned for each dimension.

This factor level contributes one parameter, *β*_0_, to the model.

**Figure 7.**
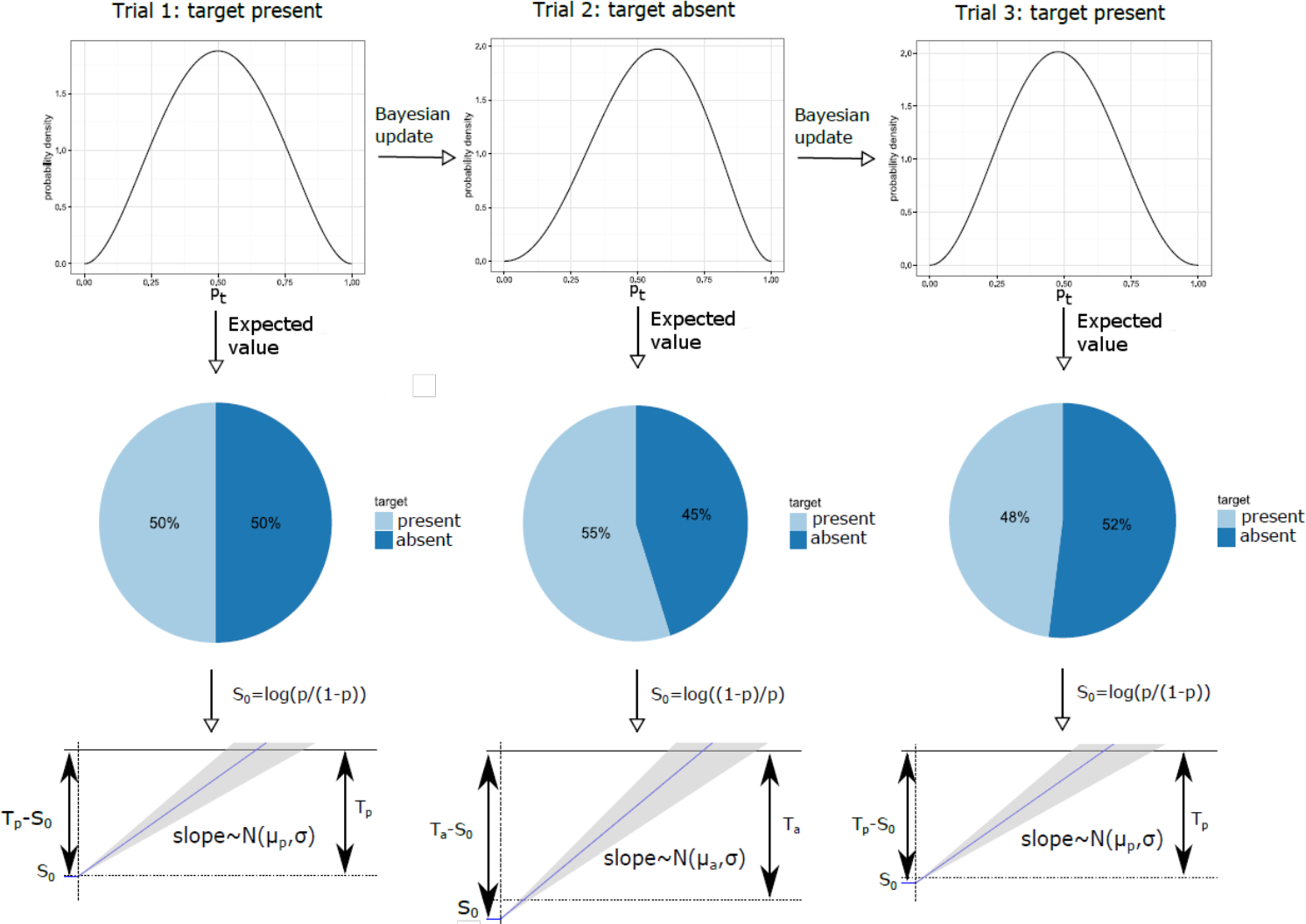
Schematic illustration of prior updating and the resulting changes of the starting point. The top panels show the hyperprior, i.e., the probability distribution on the frequency of target present trials (p), and how it changes over three subsequent trials. The middle panels show the current best estimate of the frequency distribution over target-present and -absent trials (i.e., p and 1 — p). The best estimate of p is defined as the expected value of the hyperprior. The bottom panels show a sketch of the evidence accumulation process where the starting point is set as the log prior odds for the two response options (target-present vs.-absent), computed based on the current best estimate of p. T_p_ and T_a_ are the decision thresholds for target-present and -absent responses, respectively, and and μ_p_ and μ_a_ are the respective drift rates. The sketch of the evidence accumulation process is based on the LATER model (rather than the DDM) and therefore shown with a single boundary (that associated with the correct response). Note that the boundary depicted for trial 2 (target absent) is not the same as those for (target-present trials) trials 1 and 3. In the equivalent figure based on the DDM, there would have been two boundaries, and on trial 2, the drift rate would have been negative and the starting point would have been closer to the upper boundary than on the first trial. Note also that this figure illustrates updating with some memory decay (see level 3). Without memory decay, the distribution on trial 3 would be exactly the same as on trial 1.

**Level 3** (*S*_0_ **with decay)**: Like at Level 2, *S*_0_ is updated based on the history of the RDF/TDD through Bayesian updating of the prior. In addition, we incorporated a forgetting mechanism based on the Dynamic Belief Model (DBM) [29]. That is, in addition to Bayesian updating of the probability distribution on the prior *H*_*t*_, there was, on each trial, a probability *α* with which the prior was redrawn from the starting distribution *H*_0_. This forgetting mechanism was implemented through the following equation:

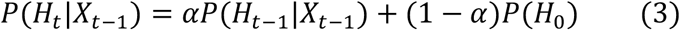

This model is identical to the fixed no-updating model (Level 1) when *α* equals 0, and is identical to the model specified in Level 2 when *α* equals 1. For intermediate values of α, the prior is partially reset to the initial prior on each trial. This factor level contributes two parameters, *α* and *β*, to the model.

For factor levels 4-6, it is the evidence accumulation rate (*r*), rather than the starting point (*S*_0_), that is being updated from trial to trial. Updating could be based on either the RDF or the TDD (in Experiment 2, these were the same), which we will refer to as the update variable (UV). In each case, UV can have two possible values, *u*_1_ and *u*_2_, namely, either color and orientation or target-present and -absent, depending on which experiment is being modelled.

**Level 4 (Binary rate)**: The RDF/TDD repetition/switch updates the information accumulation rate r in a step-wise manner, with the rate depending only on one-trial-back changes of UV: the rate is scaled by a parameter *κ*, whose value was either *κ*_0_ (0<*κ*_0_<1) when the UV changed between trials, or 1 when the UV repeated:

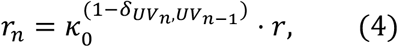

where 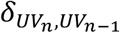 is the Kronecker delta function. When updating was performed based on the target dimension, it only affected the rate on target-present trials that were immediately preceded by a (target-present) trial with a target defined in a different dimension. This factor level contributes one parameter, *κ*, to the model.

Levels 5-6 were both designed to reduce the evidence accumulation rate after a UV switch, just like factor Level 4, but allowing for an influence from more than one trial back.

**Level 5 (Rate with decay)**: The RDF/TDD repetition/switch updates the rate *r* with a memory decay, which was accomplished by reducing the rate whenever the (value of the) UV switched between trials, and increasing it when the UV repeated. Specifically, the rate was scaled by *κ* on each trial if updating was based on the RDF, or on each target-present trial if it was based on the target-defining dimension. The starting value of *κ* was set to 1, and it was increased by Δ after each UV repetition, and decreased by Δ after each UV switch. There was also a forgetting mechanism, the same as that implemented at Level 3, such that trials further back had less influence:

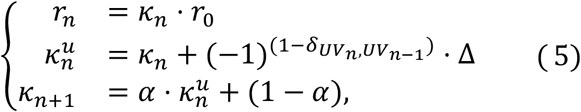

where *κ*_*n*+1_ determines the amount of scaling of the rate on trial n+1 while 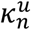 is the value of *κ* after being updated based on the stimulus on trial n, and 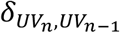 the Kronecker delta function. When updating was based on the target-defining dimension, no increase or decrease by Δ occurred on target-absent trials, while the forgetting step was still performed. This factor level contributes two parameters, Δ and α, to the model.

**Level 6 (Weighted rate)**: The RDF/TDD repetition/switch updates the rate *r* with a shared weight resource. Level 6, like Level 5, allowed for an influence on the rate from more than one trial back. Like at Levels 4 and 5, a separate rate was used for each value of the UV (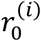 for UV = *u*_*i*_, *i* = {1,2}). Just like at Levels 4 and 5, these rates were scaled based on trial history. However, unlike Levels 4 and 5, the factors by which the two rates were scaled summed to a constant value, as if there was a shared ‘weight’ resource. After a trial on which a given value of the UV had occurred, some weight was moved to the scaling factor associated with that value of the UV (i.e., the target dimension or the target-present/absent status depending on whether the rule was used for TDD- or RDF-based updating). This updating rule was inspired by the dimension-weighting account [6].Specifically, the rate 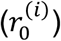 was scaled by *κ*^(*i*)^, where the summation of the scaling factor was kept constant at 2, that is,

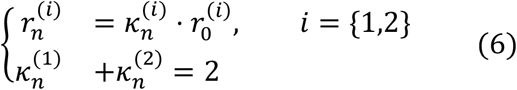

where the scaling factor 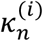, *i* = {1,2}, updates with the following rules,

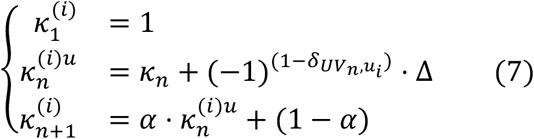

Thus, after each trial, some amount of the limited resource determining the scaling of the rate was moved to the scaling factor associated with the value of the UV that had occurred on that trial. In addition, the same forgetting rule as that implemented at Level 5 was used. When the updating was based on the target dimension, no scaling of the rate or updating of *κ* was performed on target-absent trials, though the forgetting rule was still applied, just like at Level 5.

This level contributes two parameters, Δ and *α*, to the model.

### Model comparison

With the full combination of the four factors, there were 144 (2 × 2 × 6 × 6) models altogether for comparison: non-decision time (with/without), evidence accumulation models (DDM vs. LATER), RDF-based updating (6 factor levels), and TDD-based updating (6 factor levels). We fitted all models to individual-participant data across the three experiments, which, with 12 participants per experiment, yielded 5184 fitted models. Several data sets could not be fitted with the full memory version of the starting point updating level (i.e., Level 2) of the dimension-based updating factor, due to the parameter updating to an extreme. We therefore excluded this level from further comparison.

#### Experiment 1: Target detection with variable ratios of target-present vs. -absent trials

Figures 8-10 shows the mean relative Akaike Information Criteria (AICs) for each of our experiments. For each individual participant we found the best (lowest AIC) model, then we subtracted the AIC of that model from the AIC for every model for that participant, and finally we averaged this relative AIC across all participants. In Figure 8 the mean relative AIC is shown for all models with a non-decision time component in Experiment 1 (recall that the task in Experiment 1 was to discern whether a target was present or absent; the ratio of target-present/absent trials was varied between blocks, and the target dimension, color or orientation, changed only between shorter mini-blocks). The AIC is a measure of the quality of a model, taking into account goodness of fit (as measured by the likelihood) and penalizing models with more free parameters, where lower AIC values indicate better model performance. The mean relative AIC would be zero for a model if that model was the best model for every participant; larger values indicate how much worse, on average across participants, a given model performed compared to the best model. In this figure, as well as in Figures 9 and 10 (Experiments 2 and 3), only models with a non-decision time component have been included since these generally performed better, in AIC terms, than models without a non-decision time (see Table S1 in Supplement S3). This was particularly the case when the DDM was used for RT distribution modeling (and to a lesser extent with the LATER model) – though, for each experiment, the model that achieved the lowest AIC did include a non-decision time component, regardless of whether the LATER or the DDM was used. In general, models using LATER for the RT distribution outperformed those using DDM. Of note, though, the pattern across the other factors was very similar; for instance, for the models with the lowest AIC-, the (other) factor levels were the same whether the DDM or the LATER model was used (see also Supplement S3 for figures of the AICs for the models without a non-decision time component).

**Figure 8.**
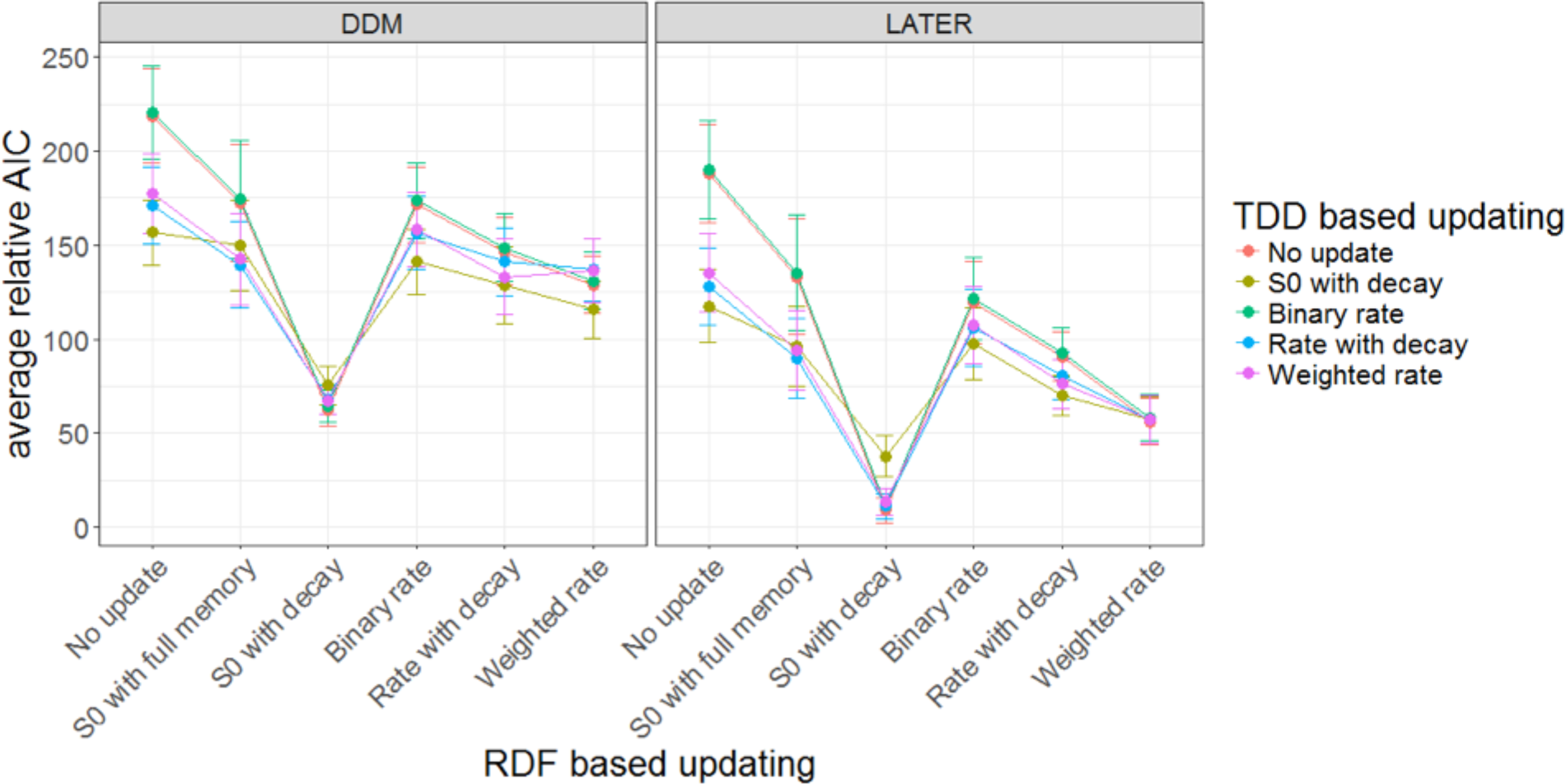
Mean relative AICs as a function of the tested models in Experiment 1. For each participant, the AIC of the best-performing model has been subtracted from the AIC for every model, before averaging across participants. Error bars indicate the standard error of the mean. The response-based updating rules are mapped onto the x-axis (RDF-based updating), while the dimension-based updating rules are indicated by different colors (TDD-based updating). The left-hand panel presents the results for the DDM, the right-hand panel for the LATER model. Only models with a non-decision time component are included in the figure. Models without a nondecision time component generally performed worse, and the best-fitting model included a nondecision time component (see also Table S2 in Supplement S4).

Importantly, in Experiment 1, for target presence/absence switches/repetitions, which (in Experiment 1) were equivalent to response switches/repetitions, the best-fitting model turned out to be that which updates the initial starting point with partial forgetting. For the dimension switch/repetition, by contrast, the various updating rules yielded comparable results, though no other rule was better than the no-update rule. The latter is unsurprising given that, in Experiment 1, the dimensions were separated in different mini-blocks, that is, effectively there was no dimension switch condition (except for the infrequent changes between mini-blocks).

#### Experiment 2: Dimension discrimination with variable ratios of color vs. orientation targets

Figure 9 depicts the mean relative AICs, averaged across all participants, for all models with a non-decision time component in Experiment 2, in which there was a target present on each trial and the task was to report the dimension of the target, color versus orientation, which changed randomly from trial to trial, and the ratio of color to orientation target trials was varied between blocks. Similar to Experiment 1, models using LATER did overall better than those using DDM. The best factor level for response-based updating involved updating of the initial starting point with partial forgetting. And the best factor level for updating based on the target dimension turned out to be updating of the accumulation rate with partial forgetting (i.e., Level 5, “rate with decay”, of the dimension-based updating factor).

**Figure 9.**
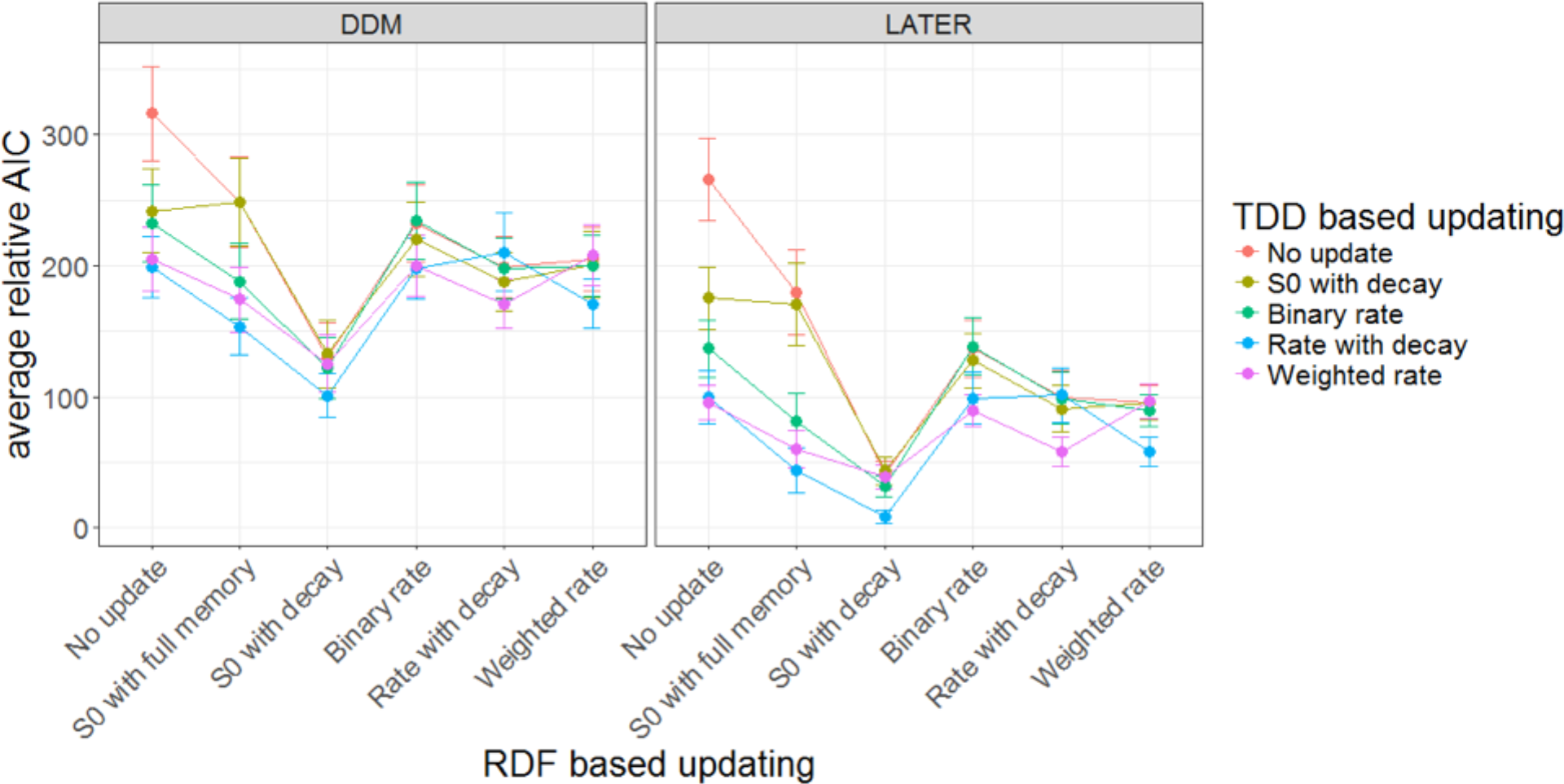
Mean relative AICs as a function of the tested models in Experiment 2. For each participant, the AIC of the best-performing model has been subtracted from the AIC for every model, before averaging across participants. Error bars indicate the standard error of the mean. The response-based updating rules are mapped onto the x-axis (RDF-based updating), while the dimension-based updating rules are indicated by different colors (TDD-based updating). The left-hand panel presents the results for the DDM, the right-hand panel for the LATER model. Only models with a non-decision time component are included in the figure. Models without a nondecision time component generally performed worse, and the best-fitting model included a nondecision time component (see also Table S2 in Supplement S4).

#### Experiment 3: Standard pop-out search task with equal target-present vs. -absent trials

Experiment 3 used a standard pop-out search detection task (target-present vs. -absent response), with color and orientation targets (on target-present trials) randomly mixed within blocks. Like Experiments 1 and 2, the LATER model and the response-based updating of the initial starting point outperformed the other model variants (see Figure 10). For dimension switches/repetitions, again a form of accumulation rate updating won over the other factor levels. The top two models both involved rate updating, with a slightly superior AIC score for the model implementing a weighting mechanism with a memory of more than one trial back (‘Weighted rate’) compared to the model in which the rate updating was based only on whether the dimension was repeated/switched compared to the previous trial (‘binary rate’).

To summarize: For all three experiments, the best models, in AIC terms, were based on the LATER rather than the DDM and used updating of the starting point with partial forgetting based on the response. For the two experiments in which color and orientation targets were randomly interleaved within each block, that is, in which dimension switching occurred, the best model involved updating of the evidence accumulation rate based on the dimension. A complementary analysis based on individual participants’ fits (Supplement S4) supports the same conclusions.

**Figure 10.**
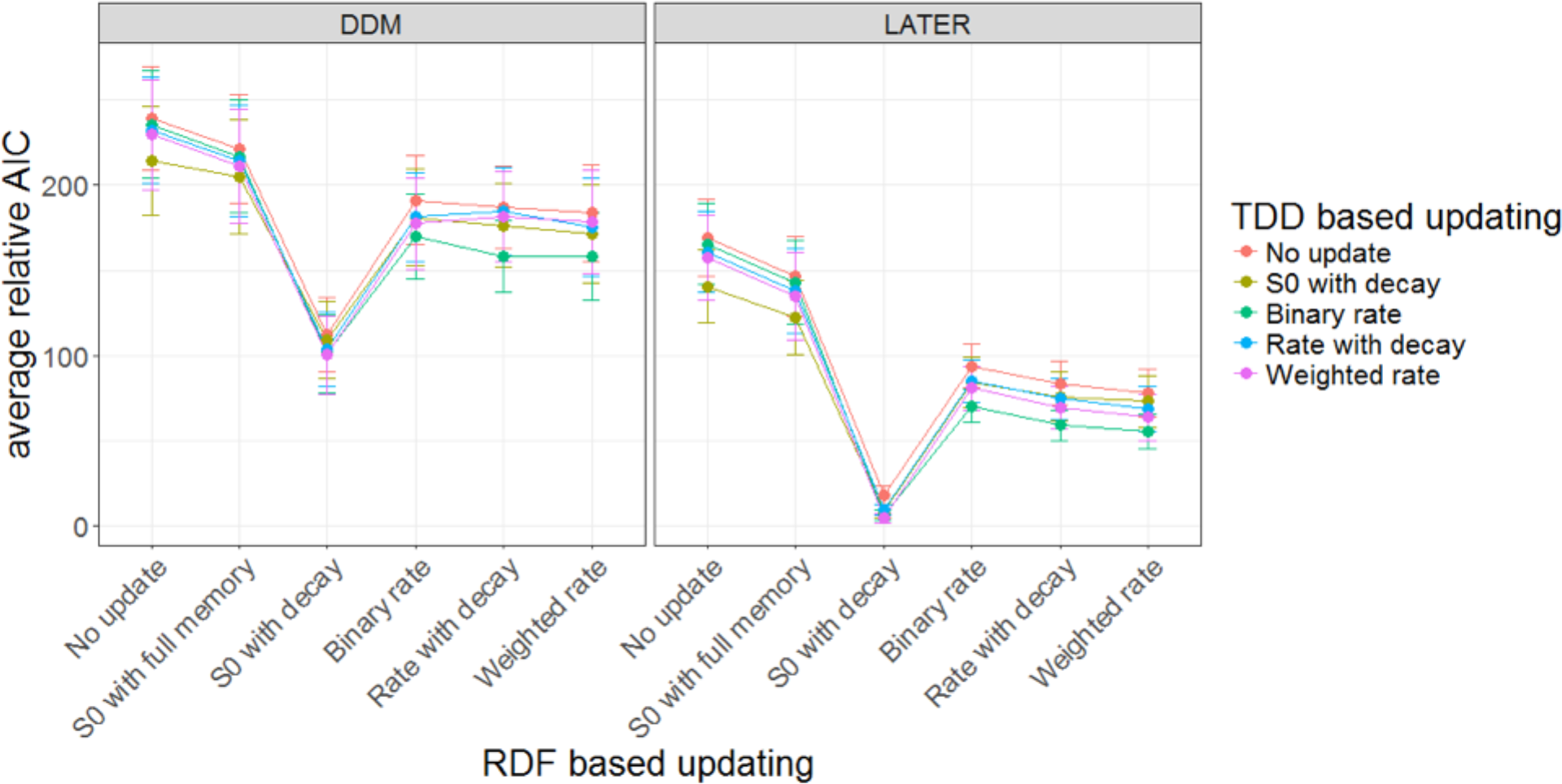
Mean relative AlCs as a function of the tested models in Experiment 3. For each participant, the AIC of the best-performing model has been subtracted from the AIC for every model, before averaging across participants. Error bars indicate the standard error of the mean. The response-based updating rules are mapped onto the x-axis (RDF-based updating), while the dimension-based updating rules are indicated by different colors (TDD-based updating). The left-hand panel presents the results for the DDM, the right-hand panel for the LATER model. Only models with a non-decision time component are included in the figure. Models without a nondecision time component generally performed worse, and the best-fitting model included a nondecision time component (see also Table S2 in Supplement S4).

### Prediction of RTs and model parameter changes

To obtain a better picture of the best model predictions, we plotted predicted versus observed RTs in Figure 11. Each point represents the average RT over all trials from one ratio condition, one trial condition, and one inter-trial condition in a single participant. There are 144 points each for Experiments 1 and 2 (12 participants × 3 ratios × 2 trial conditions × 2 inter-trial conditions) and 108 for Experiment 3 (12 participants × 3 trial conditions × 3 inter-trial conditions). The predictions were made based on the best model for each experiment, in terms of the average AIC (see Figures 8, 9, and 10). The *r*^2^ value of the best linear fit is 0.85 for Experiment 1, 0.86 for Experiment 2, and 0.98 for Experiment 3, and 0.89 for all the data combined.

**Figure 11.**
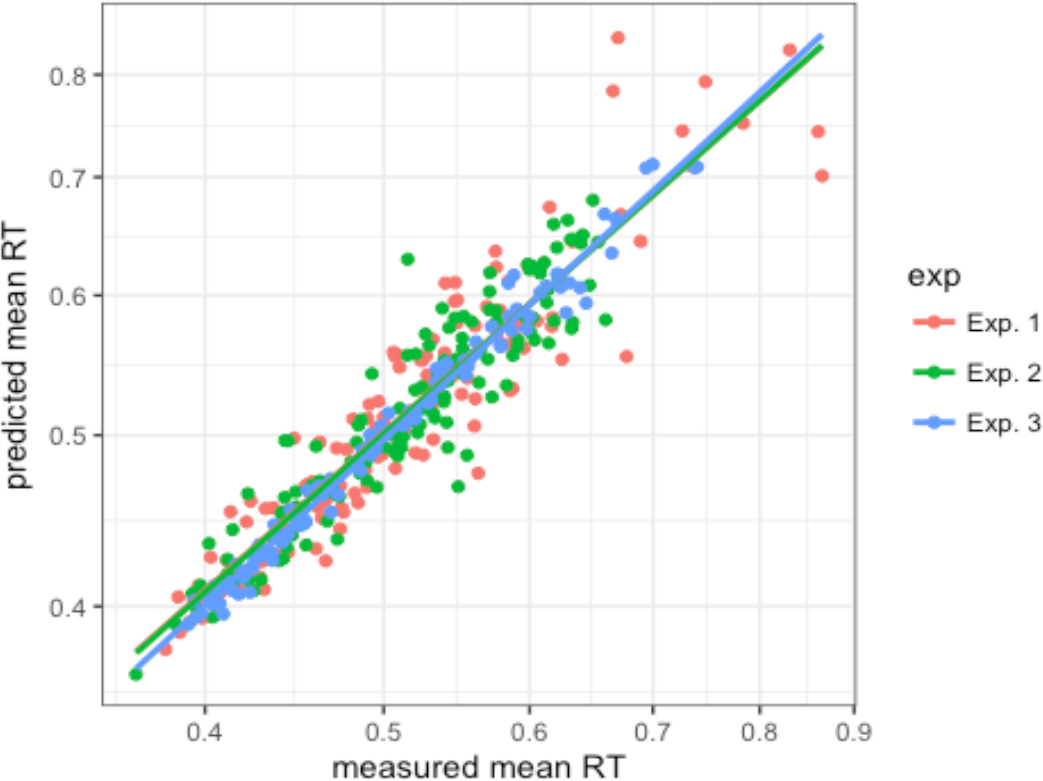
Scatterplot of predicted vs. observed mean RTs for all experiments, participants, ratio conditions, and inter-trial conditions, for each experiment. Lines show the corresponding linear fits.

Figure 12 presents examples of how the starting point (*S*_0_) and rate were updated according to the best model (in AIC terms) for each experiment. For all experiments, the best model used starting point updating based on the response-defining feature (Figure 12A, C, E, left panels). In Experiments 1 and 2, the trial samples shown were taken from blocks with an unequal ratio; so, for the starting point, the updating results are biased towards the (correct) response on the most frequent type of trial (Figure 12A, C). In Experiment 3, the ratio was equal; so, while the starting point exhibits a small bias on most trials (Figure 12E), it is equally often biased towards either response. Since, in a block with unequal ratio, the starting point becomes biased towards the most frequent response, the model predicts that the average starting point to boundary separation for each response will be smaller in blocks in which that response is more frequent. This predicts that RTs to a stimulus requiring a particular response should become faster with increasing frequency of that stimulus in the block, which is what we observed in our behavioral data. In addition, since, after each trial, the updating rule moves the starting point towards the boundary associated with the response on that trial, the separation between the starting point and the boundary will be smaller on trials on which the same response was required on the previous trial, compared to a response switch. This predicts faster RTs when the same response is repeated, in line with the pattern in the behavioral data. The forgetting mechanism used in the best models ensures that such inter-trial effects will occur even after a long history of previous updates.

In Experiment 1, the best model did not use any updating of the drift rate, but a different rate was used for each dimension and for target-absent trials (Figure 12B). In Experiment the best model updated the rate based on the ‘Rate with decay’ rule described above. The rate is increased when the target-defining dimension is repeated, and decreased when the dimension switches, across trials, and these changes can build up over repetitions/switches, though with some memory decay (Figure 12D). Since the target dimension was (also) the response-defining feature in Experiment 2, the rate updating would contribute to the ‘response-based’ inter-trial effects. In Experiment 3, the best model involved the ‘Weighted rate’ rule. Note that the rate tends to be below the baseline level (dashed lines) after switching from the other dimension, but grows larger when the same dimension is repeated (Figure 12F). This predicts faster RTs after a dimension repetition compared to a switch, which is what we observed in the behavioral data.

**Figure 12.**
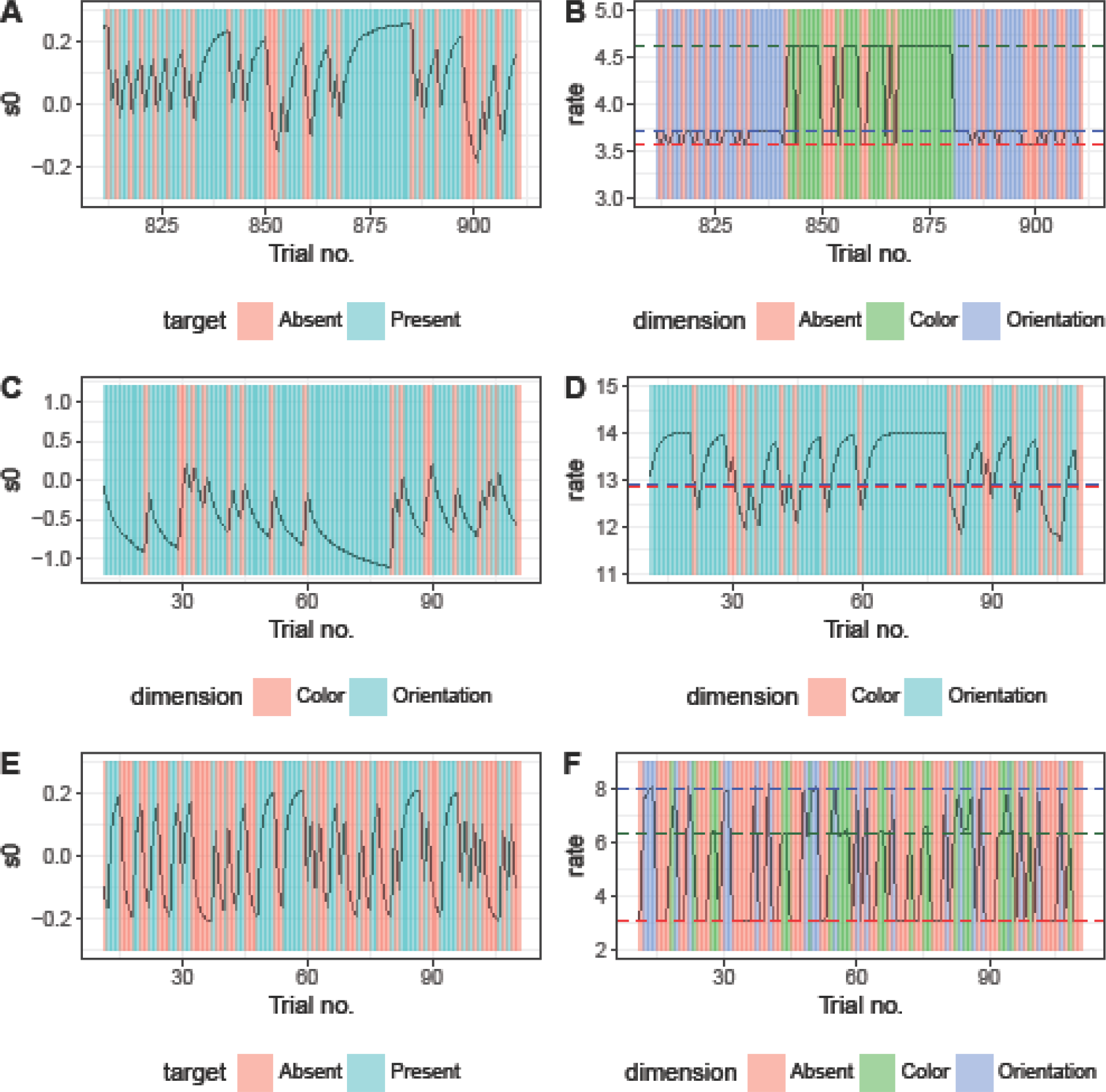
Examples of the updating of the starting point (s0) and the rate. Left panels A, C, and E show examples of starting point updating for a representative sample of trials from typical participants from Experiments 1-3. Panels B, D, and F show updating of the rate for the same trial samples (from the same participants); the dashed lines represent the baseline rates before scaling for target-absent, color target, and orientation target trials (i.e., the rate that would be used on every trial of that type if there was no updating). In each case, updating was based on the best model, in terms of average AIC, for that experiment.

## Discussion

In three experiments, we varied the frequency distribution over the response-defining feature (RDF) of the stimulus in a visual pop-out search task, that is, target presence versus target absence (Experiments 1 and 3) or, respectively, the dimension, color versus orientation, along which the target differed from the distractors (Experiment 2). In both cases, RTs were overall faster to stimuli of that particular response-defining feature that occurred with higher frequency within a given trial block. There were also systematic intertrial ‘history’ effects: RTs were faster both when the response-defining feature and when the target-defining dimension repeated across trials, compared to when either of these changed. Our results thus replicate previous findings of dimension repetition/switch effects [6,9].

In contrast to studies on ‘priming of pop-out’ (PoP) [3,32–34], we did not find significant feature-based repetition/switch effects (consistent with [6]), except for Experiment 2 in which the target dimension was also the response-defining feature. The dimension repetition/switch effects that we observed were also not as ‘long-term’ compared to PoP studies, where significant feature ‘priming’ effects emerged from as far as eight trials back from the current trial. There are (at least) two differences between the present study and the PoP paradigms, which likely contributed to these differential effect patterns. First, we employed dense search displays (with a total of 39 items, maximizing local target-to-non-target feature contrast), whereas PoP studies typically use much sparser displays (e.g., in the ‘prototypical’ design of Maljkovic & Nakayama [3,32–34], 3 widely spaced items: one target and two distractors). Second, the features of our distractors remained constant, whereas in PoP studies the search-critical features of the target and the distractors are typically swapped randomly across trials. There is evidence indicating that, in the latter displays, the target is actually not the first item attended on a significant proportion of trials (according to [35], on some 20% up to 70%), introducing an element of serial scanning especially on feature swap trials on which there is a tendency for attention (and the eye) to be deployed to a distractor that happens to have the same (color) feature as the target on the previous trial (for eye movement evidence, see, e.g., [36,37]). Given this happens frequently, feature checking would become necessary to ensure that it is the (odd-one-out) target item that is attended and responded to, rather than one of the distractors. As a result, feature-specific effects would come to the fore, whereas these would play only a minor role when the target can be reliably found based on strong (local) feature contrast [38]. For this reason, we opted to start our modeling work with designs that, at least in our hand, optimize pop-out (see also [39]), focusing on simple target detection and ‘non-compound’ discrimination tasks in the first instance. Another difference is that we used simple detection and ‘non-compound’ discrimination tasks in our experiments, while PoP experiments typically employ ‘compound’ tasks, in which the response-defining feature is independent of the target-defining feature. We do not believe that the latter difference is critical, as reliable dimension repetition/change effects have also been observed with compound-search tasks (e.g., [40]), even though, in terms of the final RTs, these are weaker compared to simple response tasks because they are subject to complex interactions arising at a post-selective processing stage (see below and [41,42]).

To better understand the basis of the effects we obtained, we analyzed the shape of the RT distributions, using the modified LATER model [26] and the DDM [21,22]. Importantly, in addition to fitting these models to the RT distribution across trials, we systematically compared and contrasted different rules of how two key parameters of the LATER/DDM models – the starting point (*S*_0_) or the rate (*r*) of the evidence accumulation process – might be dynamically adapted, or updated, based on trial history. We assumed two aspects of the stimuli to be potentially relevant for updating the evidence accumulation parameters: the response-defining feature (RDF) and the target-defining dimension (TDD; in Experiment 2, RDF and TDD were identical). Thus, in our full factorial model comparison, trial-by-trial updating was based on either the response-defining feature or the target dimension (factor 1), combined with updating of either the starting point or the rate of evidence accumulation (factor 2), with a number of different possible updating rules for each of these (6 factor levels each). An additional factor (factor 3) in our model comparison was the evidence accumulation model used to predict RT distributions: either the DDM or the LATER model. Finally, to compare the DDM and LATER models on as equal terms as possible, we modified the original LATER model by adding a non-decision time component. Thus, the fourth and final factor concerned whether a non-decision time component was used or whether the non-decision time was fixed to zero.

Our model assumes that the starting point (*S*_0_) is updated based on the observer’s current estimate of the probabilities of the response alternatives, which may depend on trial history. The assumption that the starting point is set based on the prior probabilities of the two alternative responses is consistent with a Bayesian framework of evidence accumulation, in which evidence is accumulated from the starting log prior odds until a threshold level is reached on the posterior odds before a decision is made [19,26,43]. Our model assumes that the relative frequency of the two alternative values of the RDF (target-present vs. -absent in Experiments 1 and 3, color vs. orientation target in Experiment 2) is learned from trial history. Since there is always some uncertainty about the frequency, the range of plausible values, given the trial history, is represented by a probability distribution. On the first trial, this distribution is set to a Bernoulli distribution, with a single parameter representing a prior belief about how frequently the two values of the RDF will occur before encountering the first search display. This probability distribution is then updated according to Bayes’ rule on each trial. Note that, on its own, such Bayesian updating would converge on a stable estimate and then not change much – which would predict the size of the inter-trial effects to decrease over the course of an experiment. However, we did not observe such a decrease in any of our experiments (see Supplement S5). For this reason, in addition to the Bayesian updating rule described above, we introduced a learning rule based on the Dynamic Belief Model [29], which assumes there is some fixed probability on each trial that the stimulus frequencies will change and which therefore, in addition to the Bayesian updating, involves a ‘forgetting’ step that serves to reduce the weight of old information relative to the most recent one. This model allows for rapid adaptation to a change even after a long period without any change; and, importantly, it does not predict a decrease of the inter-trial effect magnitude over the course of an experiment.

Considering the data from each experiment individually, we found that the best model (with the lowest AIC) used updating of the starting point, with partial forgetting (i.e., the learning rule from the DBM), based on the history of the response-defining feature of the stimulus array. This updating can explain both the effect of RDF frequency on RTs and the response-based inter-trial effects. The updating would result in the starting point being, on average, closer to the threshold associated with the most frequently required response in each trial block, predicting the effect of frequency on RTs. And response-based intertrial effects arise in the model because, after each trial, the starting point is moved closer to the threshold associated with the response that was required on that trial, reducing the starting point to boundary separation if that response is again required on the next trial. The forgetting mechanism ensures that the magnitude of the starting point shifts, and therefore the predicted inter-trial effects, do not shrink towards zero over the course of the 1000 plus trials in our experiments (in line with our data, which revealed no evidence of such a shrinkage; see Supplement S5). Some form of forgetting mechanism is likely to be important for adapting to a changing environment [29].

It might be argued that the frequency effects and response-based inter-trial effects on the mean RTs might, potentially, be equally well explained by trial-to-trial adaptations of the rate of evidence accumulation. However, this would have predicted a different RT distribution, and our model comparison did not favor models in which the rate was updated based on response history. We therefore conclude that the most likely explanation of response-based inter-trial effects is that observers became biased towards the response to which they assigned a higher subjective probability, and that these probabilities were particularly sensitive to what happened on the most recent trials. Of course, our starting point updating model with partial forgetting, which is closely inspired by the Dynamic Belief Model [29], is only one plausible way in which the learning of response probabilities can be implemented and linked to response biases, and other implementations remain possible. Note also that, in the present study, the feature that was critical for target detection was the same as that determining the response, which did not allow us to dissociate response repetition from target repetition effects. Further work is required to examine for such a disassociation using what is known as a ‘compound’ search task [44].

As to the dimension-based updating factor, in our model comparison, the best models differed among the three experiments. For Experiment 1, the best model did not include dimension-based updating, most likely because this experiment did not involve random dimension switching (switching occurred only between the last trial of one mini-block and the first trial of the next block, which were separated by a performance feedback screen). In Experiments 2 and 3, in which random dimension switching did occur within trial blocks, the best models involved updating of the evidence accumulation rate, though with somewhat different updating rules. For both experiments, the best model involved a rule that increased the rate when the target dimension repeated across trials and decreased it when the dimension changed. In Experiment 2, a partial memory of this increase or, respectively, decrease is then carried over to the next trial, regardless of whether the target on that trial is defined in the same or a different dimension to the preceding trial. We termed this ‘rate with decay’ rule. The best model for Experiment 3, on the other hand, used an updating rule which assumes that a different rate is associated with each dimension, where, after each trial, the rate for the dimension that defined the target on that trial is increased, and that for the other dimension is decreased by an equivalent amount. This ‘weighted rate’ rule is inspired by the dimension-weighting account [6], according to which potential target-defining dimensions share the same, limited attentional ‘weight’ resource. The two rules are similar but make significantly different predictions, for instance, when a long sequence of repeats is followed by a switch, or when a long sequence of switches occurs. The ‘rate with decay’ rule predicts the rate to be higher after a sequence of repeats followed by a single switch, compared to a switch following a run of switches – a pattern actually seen in Experiment 2 (see Supplement S6). The ‘weighed rate’ rule, by contrast, makes the opposite prediction – consistent with the pattern seen in Experiment 3 (see Supplement S6).

Recall that, in Experiment 2, the target dimension was also the response-determining feature. As a consequence, (repeatedly) switching the dimension and the response may give rise to a cost that carries over across trials by slowing the (executive) act of selecting the appropriate motor response on a given trial. This may be the case because, with choice responses, some ‘event file’ buffering the requisite S–R link might be carried over across trials and affect the speed of response decisions (see ‘episodic-retrieval theory’ below). On switch trials (‘S’), the old rule no longer applies, that is, it needs to be inhibited and replaced by a new association, where the mismatch with the old setting slows response selection. On repeated switch trials (e.g., ‘SSS’), the link relevant on the current trial (trial *n*; the same association as on trial *n*-2) might still be inhibited (from trial *n*-1, on which the rule was found to be inappropriate), slowing responses relative to switch trials preceded by repeated trials (e.g., ‘RRS’) where the association required on trial *n* is different from trial *n*-2 and would, thus not be inhibited on trial *n*−1. Assuming that the evidence accumulation in favor of a particular target dimension feeds more or less directly into the process of making a response decision, inhibition of an S-R link might narrow the whole ‘pipeline’ of perceptual and response-related evidence accumulation, explaining why the best dimension-based updating rule in Experiment 2 involved updating of the rate. This account of the cost on repeated switch trials would be consistent with the ‘negative priming’ literature (e.g., [45]).

No such cost would arise in Experiment 3, in which the dimension was not response-defining – rather, all trials with a target present (in whatever dimension it was defined) required one and the same, simple target detection response. Accordingly, dimension switches were not associated with a response switch, and so there would be no need for an updating of the S–R association after switch trials (consistent with evidence that dimensional target identity is not explicitly encoded in simple singleton detection tasks; see [9]). In this situation, on the dimension-weighting account, each repetition would mean that increasingly more weight is assigned to the repeated dimension and consequently less weight to the alternative dimension, which will be the target dimension on the switch trial at the end (RRS). Consequently, on that trial, the rate of evidence accumulation (for a target in the alternative dimension) is slowed relative to an SSS sequence (where the dimension on trial *n* had received a weight increase, rather than a decrease, on trial *n*-2). Thus, the fact the best model for that experiment involved the ‘weighted rate’ rule would lend support to ‘dimension weighting’ as the best account of dimension repetition/switch effects when there is no concurrent response switching.

Importantly, the ‘weighted rate’ and ‘rate with decay’ rules both involve updating of the rate of evidence accumulation (rather than of the starting point). The model comparison thus clearly supports the hypothesis that the dimension repetition benefit derives from more efficient stimulus processing, rather than a response bias. Convergent evidence comes from recent studies of visual search examining event-related potentials, in which dimension-specific RT inter-trial effects were reflected in the latency and amplitude of the early sensory processing N1 [46] and the N2pc component. The N2pc is commonly taken to reflect processes of spatial-attentional selection [41,47]. Thus, in light of the present model comparison, the fact that repetitions versus changes of the target-defining dimension across trials shortened the N2pc latencies would support the notion that dimension repetition increases the rate of salience accumulation for attentional target selection.

Our model comparison revealed that employing the LATER model for predicting RT distributions did a better job explaining the data than using the DDM. Note, though, that to keep the computational demands at a manageable level, we used a closed-form approximation of the RT distribution predicted by the DDM [48]. This approximation does not capture all features implemented in most computational realizations of the DDM; perhaps critically, it does not allow for trial-to-trial variability of the non-decision time. Applied to the present data, a DDM implementation with added trial-to-trial variability of the non-decision time might have significantly improved the performance of this model (whereas it would likely have made less of a difference to the LATER model) – thus reducing the difference in AIC between the LATER model and the DDM. Adding trial-to-trial variability of the non-decision time to the future model implementations may also be important theoretically, as it may be possible to explain some of this variability by adding updating rules that operate on the non-decision time. Critically though, for all the other factors in our model comparison, the best-performing levels turned out the same, whether the DDM or the LATER model was used.

Note that, while we tested a large number of possible models, there potentially are other models that might perform even better. In particular, a model that allows several updating rules to operate at once would likely perform somewhat better than our winning model. In the present study, we limited our comparisons to parsimonious models with one updating rule based on the RDF and one based on the TDD, assuming that manipulation of the RDF or the TDD only affects one distinctive process that is reflected in either the starting point *S*0 or the accumulation rate *r*. However, it remains possible that the RDF and/or the TDD influence RTs through more than one mechanism in parallel – in which case our model comparison would have identified only that mechanism which accounts for the largest portion of the inter-trial effects. In future work, it will be interesting to determine whether a model which permits the RDF and/or the TDD to operate through more than one mechanism can explain the data significantly better.

In our model, we treated target-absent trials similar to target-present trials, given that pop-out targets are detected efficiently (based on spatially parallel search), that is: with pop-out targets, a target-presence versus -absence decision can be made by setting a single threshold on the search-guiding overall-saliency map [49]. Indeed, our model predicts RTs well on both target-absent and target-present trials. However, deciding that a target is absent in a non-pop-out search task may be quite different. In a non-pop-out search display, every item in the search display would in principle need to be processed to (reliably) arrive at a correct target-absent decision [50], though some process terminating the search (and triggering a target-absent decision) prior to exhaustive scanning of all display items may also be involved [16,51]. In any case, to model non-pop-out search, a more complex model may be required in which multiple stages of evidence accumulation typically occur before a response is triggered, corresponding to checking individual items to determine whether or not they are the target.

While we examined a number of different updating rules in our model comparison, we are not suggesting that these covered all possibilities; that is, we cannot rule out that there may be updating rules that would perform even better. While our winning model was based on the Dynamic Belief Model [29], a very similar model has been proposed by Anderson and Carpenter [30], which also involves a combination of Bayesian updating and forgetting of old trials, and this could have served as an equally good starting point for our model. Another, similar model was proposed by Mozer et al. [52]. Unlike the present model, this does not involve a hyperprior on the stimulus category probability with Bayes’ rule; rather, it updates the probability more directly, using a weighted-averaging rule, with the weight assigned to older trials decaying exponentially. This rule is close to the forgetting rule of the Dynamic Belief Model. Mozer et al. [52] showed that their model can qualitatively reproduce the pattern of results from a number of ‘priming of pop-out’ and visual search experiments [3,4,52,53]. Different to our model, the model of Mozer et al. learns conditional probabilities, which they argued was essential for explaining interactions between the inter-trial effects for different features of the stimuli in some of the experiments they modeled. While learning of conditional probabilities was not necessary to explain the results from the three experiments reported here, any more complete model of inter-trial effects in visual search may well need to incorporate conditional probabilities to provide a truly general account. Another noteworthy difference to our model is that the model of Mozer et al. only included the learning of probabilities without specifying how these learned probabilities influence the perceptual decision process. Consequently, they could not make quantitative predictions about RTs and their distributions. In contrast, our model makes quantitative predictions because it combines a Bayesian updating rule with a model of the perceptual decision process (either DDM or LATER).

Another modeling framework that has previously been applied to explaining inter-trial effects in visual search is the ‘Theory of Visual Attention’ (TVA) [54]. TVA models the rate at which visual categorizations of the type “object *x* has feature *i*” are made and encoded into visual short-term memory (mediating overt responses). Each visual object receives an attentional weight, which is the product of the strength of the sensory evidence that the object belongs to category *i* and the current importance of attending to category *i*, referred to as the ‘pertinence’ of the category, summed over all relevant visual categories (i.e., categories for which there is sensory evidence). The scaling factors in our dimension-weighted rate updating rule, representing the current weight or importance assigned to each dimension, play a similar role to the pertinence values in TVA. Ásgeirsson et al. [55,56] have shown that color priming effects in visual search can be well explained by TVA, by assuming that the pertinence of a given feature increases or decreases when the target or, respectively, a distractor possesses that feature. Similarly, our dimension-weighted rate rule assumes that the scaling factor increases for a given dimension when the target is defined in this dimension, and decreases when the target is defined in a different dimension. Our finding that this was the best rule for explaining performance in Experiment 3 is thus broadly consistent with the TVA-based model proposed by Ásgeirsson et al. [48,49]. However, our model also differs from theirs in a number of respects. First, in our model, the scaling factors were associated with dimensions rather than individual features (recall that, in our paradigms, feature-specific inter-trial effects are relatively unsubstantial compared to dimension-specific effects; see also [6]). Second, the model of Ásgeirsson et al. only considered effects from a single trial back, while our dimension-weighted rate rule can model longer-term effects (of course, it would be possible to combine TVA with a similar rule to take longer-term inter-trial history into account). Third, unlike the model of Ásgeirsson et al., our model did not include ‘spatial weights’ associated with potential target locations. Ásgeirsson et al. showed that their TVA-based model performed better when taking spatial weighting into account. Note, though, that spatial weighting is likely to be more important with sparse displays and a limited set of locations (six in Ásgeirsson et al.), compared to the dense displays used in our experiments [57]. Finally, by modelling full RT distributions, we could make a distinction between two different ways in which the speed of a perceptual decision could be increased: by increasing the rate at which relevant sensory evidence accumulates or by decreasing the amount of evidence required to make a decision (through a shift of the starting point). TVA does not make any equivalent distinction.

Another framework for understanding inter-trial effects in visual search is offered by the episodic-retrieval account [14,58] – though the evidence for this account derives exclusively from compound-search tasks not investigated here. Huang et al. [4] argued that repetition effects in visual search are well explained by episodic-retrieval theory, based on the finding that repetition of a task-irrelevant feature (in their experiments: color) speeded search only when the target-defining feature (size) was also repeated (participants had to respond to the orientation of a size-defined target, irrespective of the target color). When the target-defining feature changed, RTs were slower if the task-irrelevant feature was repeated. The episodic-retrieval account can explain this pattern by assuming that participants retrieve an episodic memory trace of the target from the previous trial, which influences a post-selective process of verifying whether a candidate target is the actual target. If the retrieved memory trace completely matches the target on the current trial, the decision will be fast; by contrast, a partial match (i.e., a target of the same size but a different color) gives rise to ‘inconsistency’ and may thus be slower to process than a complete mismatch, explaining the interaction between repetition of target-defining and task-irrelevant features in the study of Huang et al. [4]. A similar result was reported by Töllner et al. [46], though for two task-relevant target attributes. They observed a partial-repetition cost when the response-defining feature (target orientation) changed across trials while the target-defining dimension (color or shape) was repeated. However, the latency of the N2pc was affected only by repetition/switch of the target-defining dimension, independently of whether the response-defining feature repeated/changed – leading Töllner et al. to conclude that at least one critical component of the target repetition/switch effect arises at a (pre-attentive) stage of saliency coding, leading up to target selection. The partial-repetition effect, by contrast, arises at a post-selective stage where the response-defining target feature is analyzed and a response decision is determined. This process is modulated by ‘linked expectancies’ between the dimension and the response: when the dimension is repeated, the system expects the response to be repeated as well, yielding a cost when the response actually changes. – Our best-fitting model, while predicting a RT cost when the dimension or the response changes (compared to when both are repeated), does not predict a larger cost when either one or the other changes, compared to when both change (instead, the dimension and response change costs would be additive). To account for such partial-repetition cost effects, further modeling work is required based on RT performance in simple-detection and compound-search tasks that make the same demands with regard to target selection, but different demands with regard to response selection (i.e., simple detection of a target-defining attribute vs. discrimination of a separate, response-defining feature), as well as RT performance in a non-search task that makes no demands on target selection, but similar demands to compound search on response selection (along the lines of [12]). RTs could then be modeled, for instance, as a series of two diffusion processes (one for target selection and one for response selection), where parameters of the second process (*r*,θ,or *S*_0_) might be set conditional upon repetition/switch of the target-defining attribute. Such a model might then also be able to account for partial-repetition costs attributable to completely (detection- and response-) irrelevant target attributes [4], over and above those caused by relevant features [46], perhaps by making updating based on irrelevant features conditional on relevant features [52].

In conclusion, we found that RTs in pop-out visual search are faster when the response required on a given trial occurred frequently in the recent past, and particularly when the same response is repeated from the previous trial. By performing a factorial model comparison, we showed that these effects are best explained by updating of the starting point of an evidence accumulation process, that is, they reflect a bias towards a response that is more likely to occur, given the recent history. We also found that reaction times are faster when the target-defining dimension is repeated, even when this is unrelated to the response. Our model comparison showed that this effect is best explained by trial-to-trial updating of the evidence accumulation rate. This suggests that dimension repetition/switch effects do not reflect a response bias, but rather reflect more efficient processing when the same dimension is repeated.

## Methods

### Experiment 1

#### Participants

Twelve subjects participated in Experiment 1 (eight females; age range 20 and 33 years). All had normal or corrected-to-normal vision and naive to the purpose of the experiment. All participants gave informed consent prior to the experiment. The study was approved by the LMU Department of Psychology Ethics Committee and conformed to the Helsinki Declaration and Guidelines.

#### Apparatus and Stimuli

Stimuli were presented on a CRT monitor (screen resolution of 1600 × 1200 pixels; refresh rate 85 Hz; display area of 39×29 cm). Participants were seated at a viewing distance of about 60 cm from the monitor. All stimuli were presented using Matlab (The Mathworks) and Psychtoolbox [59,60].

Each stimulus display consisted of 39 bars, arranged around three concentric circles (see Figure 13). The distractors were turquoise-colored vertical bars (CIE [Yxy]: 44.9, .0.23, 0.34). When a target was present, it was always on the middle circle. Targets were bars that differed from the the distractors in terms of either color or orientation, but never both. Color targets were either green (CIE [Yxy]: 45.8, 0.29, 0.57) or purple (CIE [Yxy]: 41.5, 0.29, 0.24), while orientation targets were tilted 30° clockwise or counterclockwise from the vertical. The search display subtended approximately 7.5° × 7.5° of visual angle and each individual bar had a size of approximately 0.5° × 0.1°.

**Figure 13.**
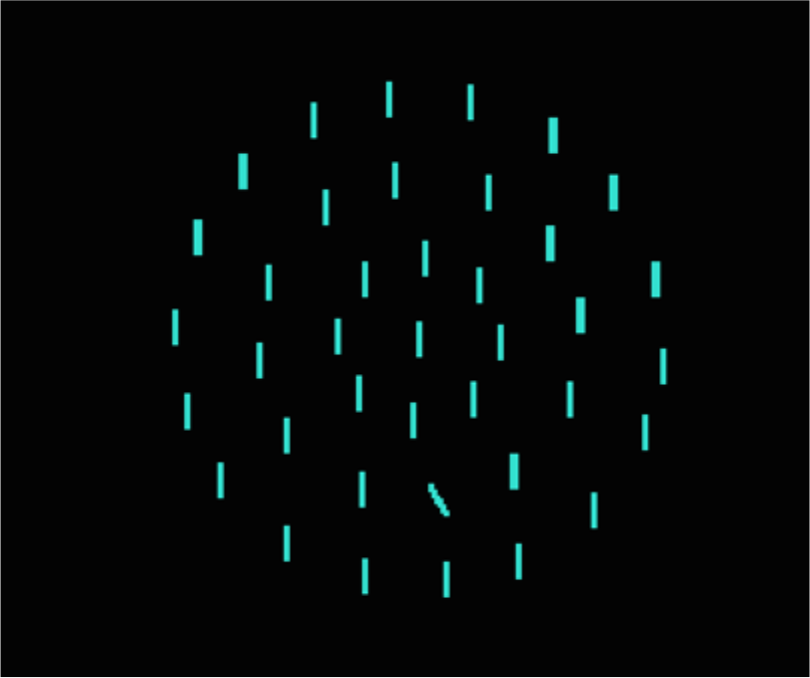
Example of visual search display with an orientation target.

#### Procedure

The experiment consisted of 30 blocks of 40 trials, divided into three equally long sections with different proportions of target-present (and, correspondingly, target-absent) trials: 75% [target-absent: 25%], 50% [50%], and 25% [75%]. A text message informed participants about the current proportion of target-present trials at the start of each block. Alternating trial blocks presented exclusively color targets or orientation targets, on target-present trials. The task was to report as quickly and accurately as possible whether a target was present or absent, using the left and right mouse buttons, respectively. Each trial started with the presentation of a fixation dot for 700-900 ms followed by the stimulus display, which was displayed until the participant responded. After the response, there was another 400-600 ms delay before the next trial started with the presentation of the fixation dot, so the total interval from response on one trial to presentation of the search display on the next trial was 1100-1500 ms.

### Experiment 2

#### Participants

Twelve new participants took part in Experiment 2 (six females; age range 18 and 33 years). All had normal or corrected-to-normal vision and were naive as to the purpose of the experiment. All participants gave informed consent before the experiment. The study was approved by the LMU Department of Psychology Ethics Committee and conformed to the Helsinki Declaration and Guidelines.

#### Apparatus and Stimuli

The same equipment and stimuli were used as in Experiment 1.

#### Procedure

The procedure was the same to Experiment 1, except that instead of reporting whether a target was present or absent, participants had to report whether the target differed from distractors in terms of color or orientation. As in Experiment 1 there were three sections, each consisting of 10 blocks of 40 trials. Unlike in Experiment 1, a target was present on every trial and it was the proportion of color (or, respectively, orientation) targets that differed between the three sections, using the same ratios of 75% [orientation: 25%], 50% [50%], and 25% [75%]. Also unlike in Experiment 1, participants were not informed in advance of what that the proportion of color trials would be in any section of the experiment, nor were they informed that this proportion would differ across the different sections of the experiment.

### Experiment 3

#### Participants

12 participants took part in Experiment 3 (six females; age range 23 and 33 years). All had normal or corrected-to-normal vision and were naive as to the purpose of the experiment. All participants gave informed consent before the experiment. The study was approved by the LMU Department of Psychology Ethics Committee and conformed to the Helsinki Declaration and Guidelines.

#### Apparatus and Stimuli

The same equipment and stimuli were used as in Experiment 1.

#### Procedure

As in Experiment 1, participants had to report on each trial whether a target was present or absent. However, the procedure differed from Experiment 1 in two important ways. First, in Experiment 3, the target-present/absent ratio was fixed at 50% throughout the whole experiment. Second, color targets and orientation targets were interleaved within each block. We used a De Bruijn sequence generator [61,62] to obtain a trial sequence where each of the four possible target types (i.e., purple, green, left-tilted, and right-tilted) were equally often followed by each target type (including itself) and were also equally often followed by a target-absent trial as by a target-present trial. Having such a trial sequence within each block requires 65 trials per block instead of 40 as in Experiments 1 and 2.

#### Bayes factors

Bayesian ANOVA and associated post-hoc tests were performed using JASP 0.86 (http://www.jasp-stats.org) with default settings. All Bayes factors for main effects and interactions in the ANOVA are ‘inclusion’ Bayes factors calculated across matched models. Inclusion Bayes factors compare models with a particular predictor to models that exclude that predictor. That is, they indicate the amount of change from prior inclusion odds (i.e., the ratio between the total prior probability for models including a predictor and the prior probability for models that do not include it) to posterior inclusion odds. We used inclusion Bayes factors calculated across matched models meaning that models that contain higher order interactions involving the predictor of interest were excluded from the set of models on which the total prior and posterior odds were based. Inclusion Bayes factors provide a measure of the extent to which the data support inclusion of a factor in the model. Bayesian t-tests were performed using the ttestBF function of the R package ‘BayesFactor’ with the default setting (rscale=“medium”).

## Modelling

To find the model that best explained our data, we performed a factorial model comparison. Full descriptions of the four factors and their levels are given in the modelling section. Here we describe the general procedure used for the model fitting, which was the same for all models.

Each model consisted of an evidence accumulation model: either the LATER model or the DDM, and two updating rules, each of which specified how one parameter of the evidence accumulation model should change from trial to trial, based on the stimulus history. There was one such updating rule for the starting point and one for the evidence accumulation rate, and in each case one of the factor levels specified that no updating at all should take place. For the DDM, we used a closed-form approximation [48], adding a scaling parameter that determined the size of the random component of the drift diffusion model. This was necessary since our rule for updating the starting point made the scale non-arbitrary.

Models were fitted using maximum likelihood, using the R function ‘constrOptim’ to find minimum value of the negative log likelihood. Error trials and outliers were excluded from the calculation of the likelihood, but were included when implementing the updating rules. Outliers were defined as trials with reaction times more than 1.5 interquartile ranges below the mean or longer than 2 seconds.

To make sure we found the best possible fit for each combination of factor levels, we used an inner and an outer optimization process. The inner optimization process was run for each combination of parameters that was tested by the outer optimization process, to find the best possible values of the inner parameters for those values of the outer parameters. The inner parameters were the parameters of the evidence accumulation model itself, except for the non-decision time which was an outer parameter (because one level of one of the factors specified that the non-decision time should be fixed to zero). For the LATER model, the inner parameters were the starting point boundary separation, and the mean and standard deviation of the distribution for the rate. For the DDM, the inner parameters were the starting point boundary separation, the rate, and the scaling parameter. These parameters could differ between target absent trials, as well as between the two different target dimensions, meaning that there were nine inner parameters for Experiments 1 and 2 and six for Experiment 2 (where there were no target absent trials). The outer parameters were the non-decision time (when this wasn’t fixed to zero), and 0 to 2 parameters for each updating rule (see the modelling section for details). This means that models could have 0 to 5 outer parameters in total depending on the factor levels.

## Supplementary Analyses

### S1. Statistical tests of error rates

Post-hoc comparisons of the error rates among blocked target frequencies revealed that fewer errors were made in high- as compared to medium- and low-frequency blocks, *t*(11) = 1.7, *p* > 0.3, *BF* = 1.2, and, respectively, *t*(11) = 3.72, *p* < 0.01,BF = 248 (Bonferoni-corrected p-value), and in medium as compared to low-frequency blocks, *t*(11) = 3.21, *p* < 0.05, *BF* = 53 (Bonferoni-corrected p-value), in Experiment 1; and similarly in high-relative to medium- and low-frequency blocks, *t*(11) = 2.07, *p* = 0.15, *BF* = 16, and, respectively, *t*(11) = 4.92, *p* < 0.001, *BF* = 228 (Bonferoni-corrected p-value), and in medium-relative to low-frequency blocks, *t*(11) = 2.85, *p* < 0.05, *BF* = 7.9 (Bonferoni-corrected p-value), in Experiment 2.

There was no interaction between target condition and frequency in either Experiment 1 or Experiment 2, *F*(2,22) = 0.83, *p* = 0.45, *BF* = 0.24, and, respectively, *F*(1.28,14.04) = 0.76 (Huynh-Feldt Corrected degrees of freedom), *p* = 0.43, *BF* = 0.29 – suggesting the effect of the frequency of a condition within a block is independent of the target stimuli.

### S2. Statistical tests of mean RTs

In Experiment 1, RTs were significantly faster in the high-frequency compared to both the low- and medium-frequency blocks [ *t*(11) = 3.96, *p* < 0.01, *BF* = 5.57 * 10^4^, and, respectively, *t*(11) = 3.88, *p* < 0.01, *BF* = 32.6 (Bonferroni-corrected p-values)], while there was no significant difference between the medium- and low-frequency blocks, [*t*(11) = 0.086, *p* > 0.9, *BF* = 0.22]. Similarly, in Experiment 2, RTs were significantly faster in the high-frequency compared to the low- and medium-frequency blocks [*t*(11) = 7.72, *p* < 0.001, *BF* = 2.78 * 10^u^, and, respectively, *t*(11) = 3.66, *p* < 0.01, *BF* = 90 (Bonferroni-corrected p-values)], and also faster in the medium-compared to the low-frequency block [*t*(11) = 4.06, *p* < 0.01, *BF* = 455 (Bonferroni-corrected p-value)].

Bayesian repeated-measures ANOVAs showed that there was no difference between trials with color- and orientation-defined targets in Experiment 2, *F* (1,11) = 0.45, *p* = 0.52, *BF* = 0.26. Interestingly, there was no interaction between target condition and frequency in either Experiment 1 or 2 [ *F*(2,22) = 2.44, *p* = 0.11, *BF* = 0.39, and, respectively, *F*(2,22) = 0.87, *p* = 0.43, *BF* = 0.31] – suggesting that the effect of the frequency is independent of the target stimuli.

### S3. Modeling results without non-decision time

Figures S1, S2, and S3 depict the mean AICs, averaged across all participants, for all models without a non-decision time (NDT) component for Experiments 1, 2, and 3 respectively. **Error! Reference source not found.** shows the differences in AIC between models without and with a non-decision time component. Compared to models with a nondecision time component (see the Modeling section in the main text), the average AICs were in general higher, that is, the models without a NDT component performed worse. Nevertheless, the dependence of the AIC on the other factors is very similar: in particular, the best combination of the other factor levels, for each experiment, is the same whether models without or with a NDT component are considered. One noticeable difference is that without a NDT component, the difference in AIC between models using the DDM and those using the LATER model is larger. Thus, while a NDT component is important for providing a good model fit, both for the DDM and for the LATER model, it appears that the LATER model can to some extent compensate for not including such a component through adjustment of the other parameters.

**Figure S1.**
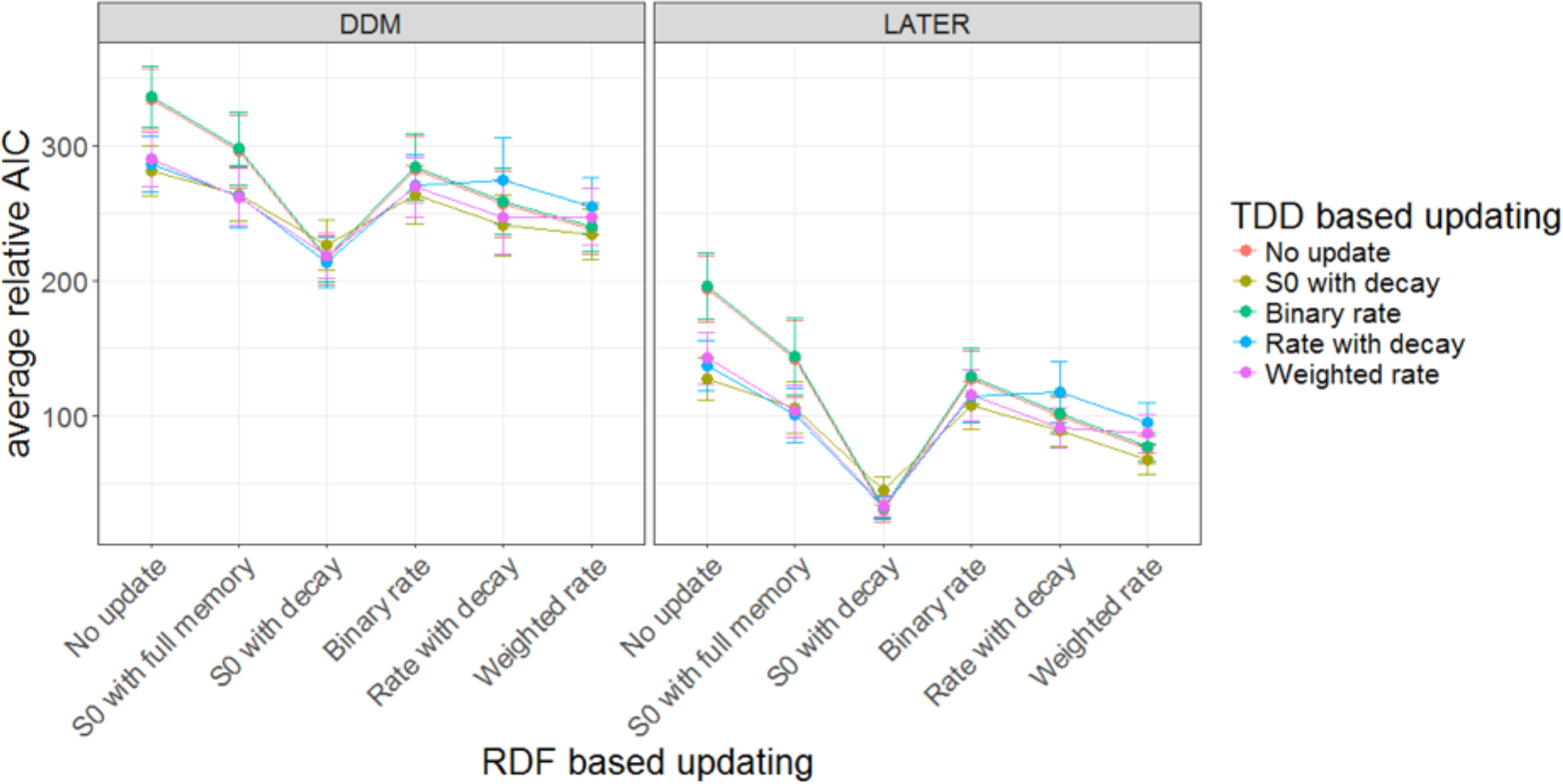
Mean relative AlCs as a function of the tested models in Experiment 1. For each participant, the AIC of the best-performing model has been subtracted from the AIC for every model, before averaging across participants. Error bars indicate the standard error of the mean. The response-based updating rules are mapped on the x-axis, the dimension-based updating rules are indicated by different colors. The left panel depicts results for the DDM, the right panel for the LATER model. Only models without a non-decision time component are included in the figure.

**Figure S2.**
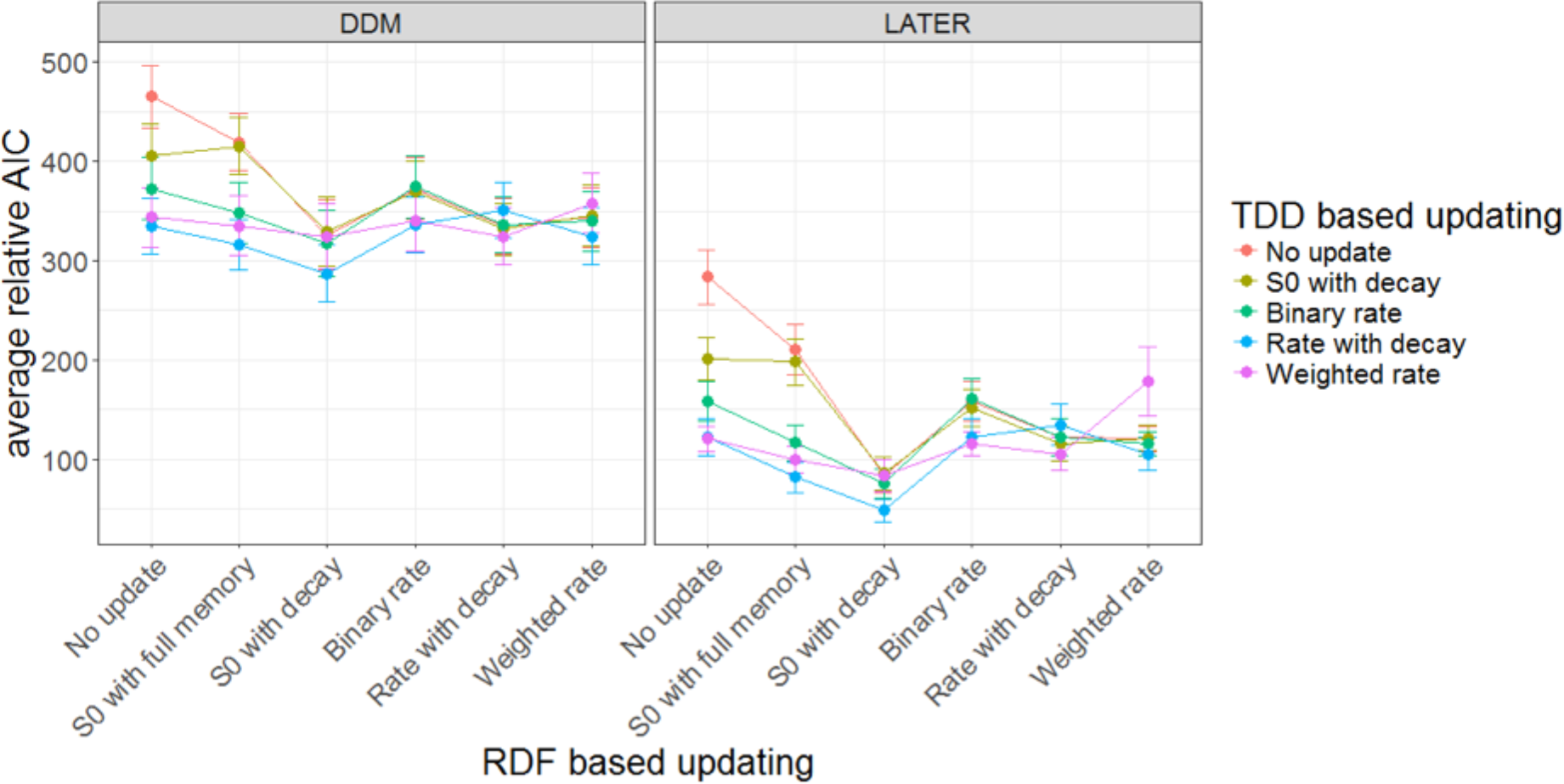
Mean relative AICs as a function of the tested models in Experiment 2. For each participant, the AIC of the best-performing model has been subtracted from the AIC for every model, before averaging across participants. Error bars indicate the standard error of the mean. The response-based updating rules are mapped on the x-axis, the dimension-based updating rules are indicated by different colors. The left panel depicts results for the DDM, the right panel for the LATER model. Only models without a non-decision time component are included in the figure.

**Figure S3.**
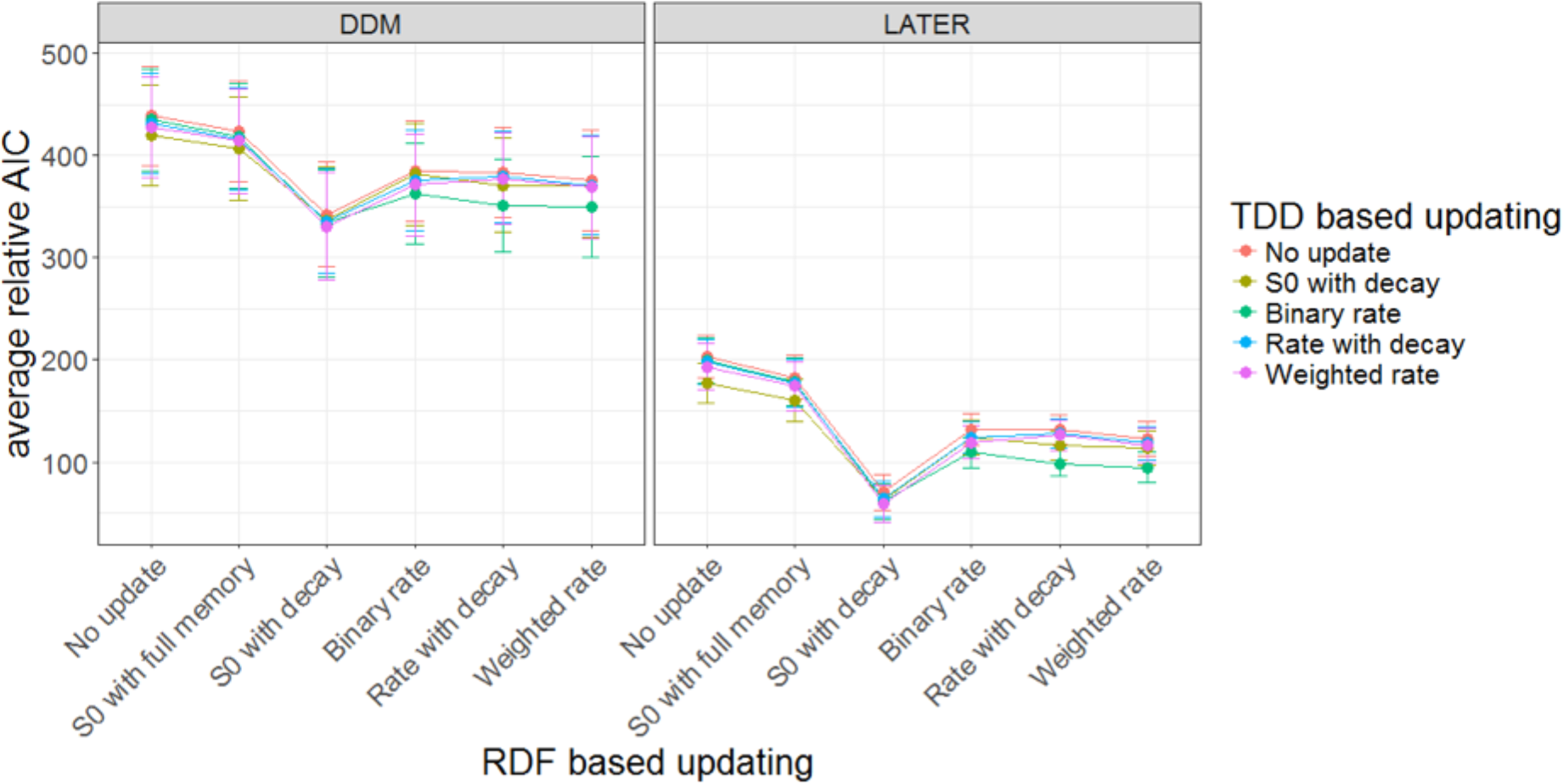
Mean relative AICs as a function of the tested models in Experiment 3. For each participant, the AIC of the best-performing model has been subtracted from the AIC for every model, before averaging across participants. Error bars indicate the standard error of the mean. The response based updating rules are mapped on the x-axis while the dimension based updating rules are mapped to different colors. The left panel contains results for the DDM while the right panel contains results for the LATER model. Only models without a non-decision time component are included in the figure.

**Table S1.**
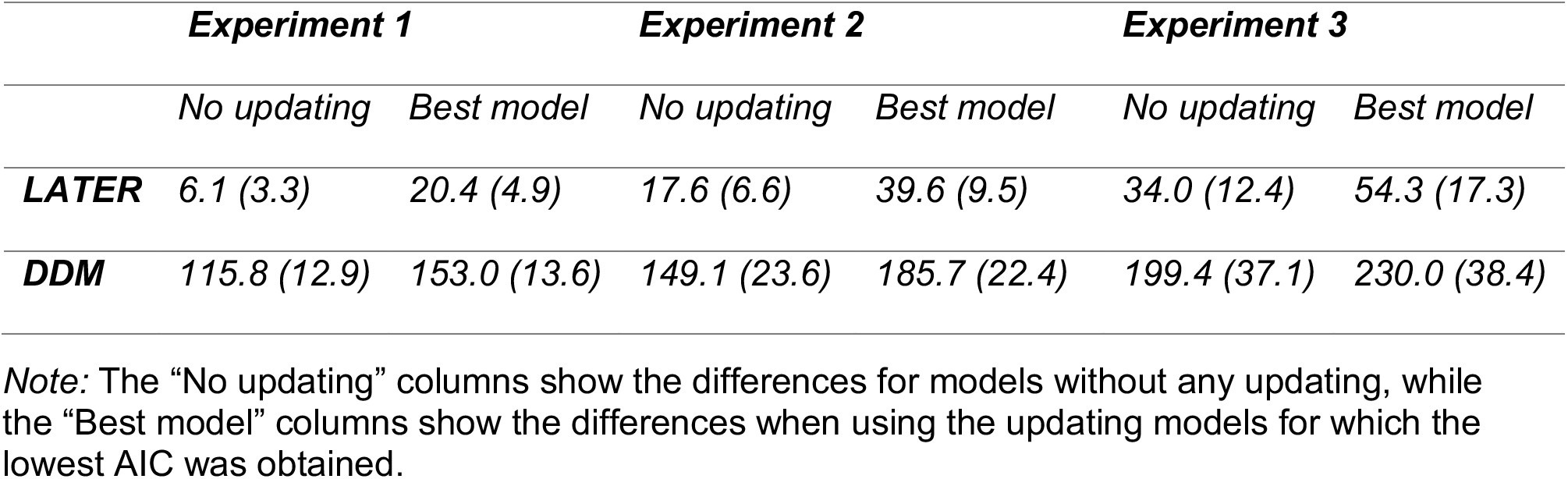
Means (and associated standard errors) of the differences in AICs between models without and with a non-decision time component.

### S4. Model comparison based on individual participants

Another way of comparing the models is by picking the best model, in terms of AIC, for each participant and counting how often each level of each factor appears in the resulting list. Table S2 summarizes the results of such an analysis. The table has four sections corresponding to the different factors of our factorial model comparison: RT distribution model, non-decision time, response-based updating, and dimension-based updating. Almost all the best-fitting models were based on the LATER model, rather than the DDM – consistent with the analyses based on average AIC values above. Also, nearly all best-fitting models included a non-decision time parameter. Inclusion of a non-decision time is, of course, unsurprising for (the best-fitting) models based on the DDM. Standard versions of the LATER, by contrast, have hardly ever included a non-decision time component. Our results suggest that adding such a component improves the fit to the data sufficiently to motivate this extra parameter.

For the response-based updating, almost all the best-fitting models were based on updating of the starting point, rather than the rate. In addition, most of these models included updating with forgetting, although for the data from Experiment 2, the model that used updating with full memory was the best-fitting model for almost as many participants. Recall that in Experiment 2, response repetitions/switches coincided with dimension repetitions/switches, so that there may have been ‘cross-talk’ between the two. The relatively good performance of the full-memory updating rule may be a consequence of such cross-talk; alternatively, it may be related to the explicit dimension (color vs. orientation) discrimination task in Experiment 2, which may have caused the memory of responses on previous trials to decay more slowly, compared to the simple-detection (target-present vs. -absent) task used in Experiments 1 and 3.

Finally, for the dimension-based updating, the best-fitting models differed among experiments. In Experiment 1, no version of dimension-based updating improved the fits sufficiently to motivate the extra parameter(s). For Experiments 2 and 3, by contrast, the various rate-based updating models most frequently provided the best account of the data. Given that the dimension only varied between mini-blocks in Experiment 1 (rather than varying randomly within each block, as in Experiments 2 and 3), it is little surprising that dimension-based updating played no significant role in modeling the data from Experiment 1. With regard to Experiment 2, it is less clear why the ‘rate with decay’ model consistently outperformed the other two rate-based updating models, while the ‘binary rate’ and ‘weighted rate’ models performed better in Experiment 3. There are two important differences between Experiments 2 and 3. First, the former used a dimension discrimination task, the later a detection task. Also, there were no target-absent trials in Experiment 2, whereas 50% of the trials were ‘target-absent’ in Experiment 3, so that dimension-based updating would affect only half of the trials. This could perhaps explain why it was worth the extra parameter to have a longer memory than a single trial back for dimension-based updating in Experiment 2, but not in Experiment 3 (where the ‘binary rate’ model, with a memory of only a single trial back but one less parameter, most frequently provided the best fit to the data). The better performance of the ‘Rate with decay’, compared to the ‘weighted rate’, version of rate updating in Experiment 2 is harder to explain with the present design; it may well have to do with the use of a discrimination (rather than a detection) task or the absence of no-target trials, but this requires further investigation.

Overall, the results of this form of model comparison closely matched those of comparing models based on the average AIC values: the factor levels that show up most frequently in the list of the best-fitting models are identical to those with the lowest average AICs, with just one exception: the ‘binary rate’ level would be preferred over the ‘weighted rate’ for dimension-based updating in Experiment 3. Importantly both the ‘binary rate’ and ‘weighted rate’ updating rules involve updating of the evidence accumulation rate, even though they differ in that the ‘weighted rate’ rule has a memory of more than one trial back.

**Table S2.**
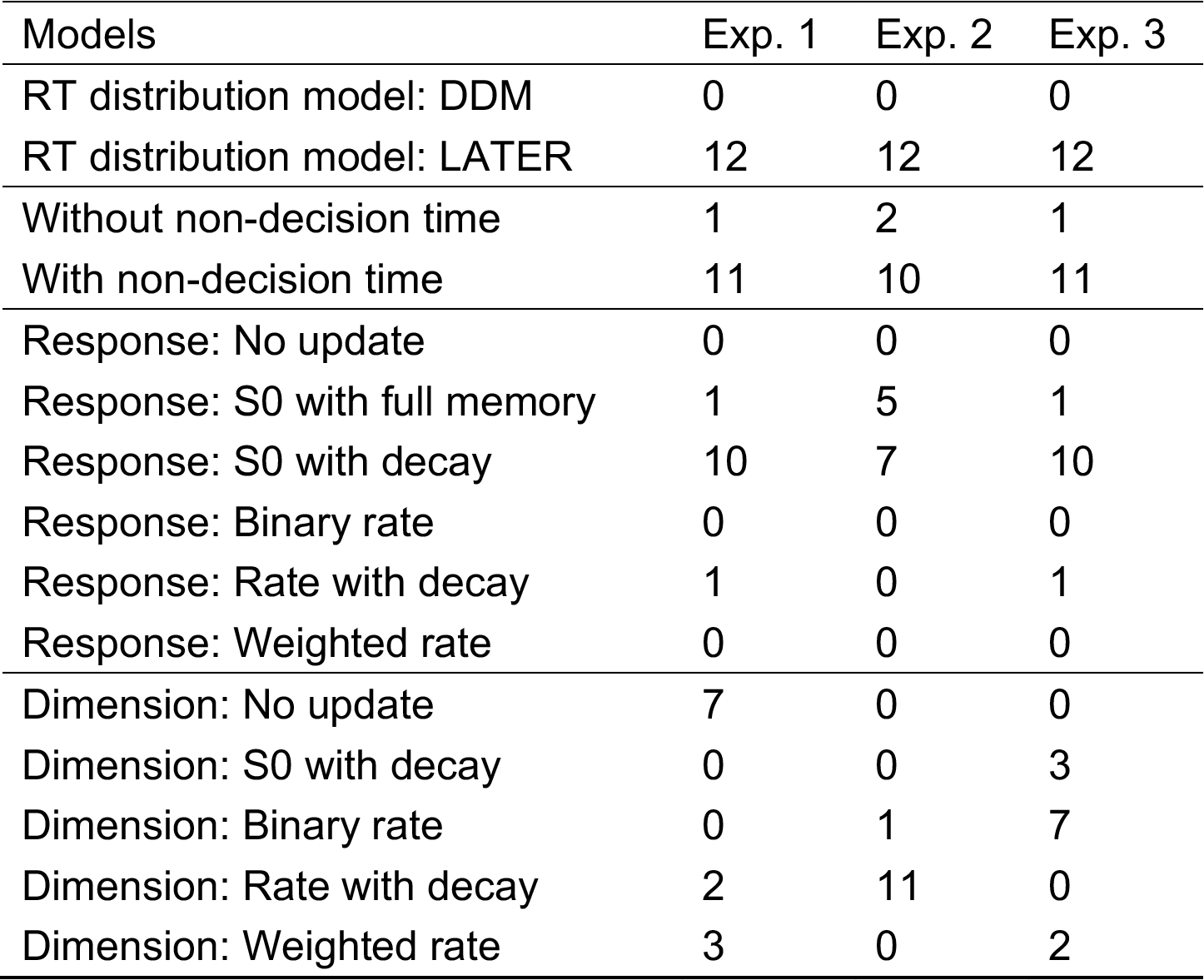
Model comparison across individual participants

| Models | Exp. 1 | Exp. 2 | Exp. 3 |
| --- | --- | --- | --- |
| RT distribution model: DDM | 0 | 0 | 0 |
| RT distribution model: LATER | 12 | 12 | 12 |
| Without non-decision time | 1 | 2 | 1 |
| With non-decision time | 11 | 10 | 11 |
| Response: No update | 0 | 0 | 0 |
| Response: S0 with full memory | 1 | 5 | 1 |
| Response: S0 with decay | 10 | 7 | 10 |
| Response: Binary rate | 0 | 0 | 0 |
| Response: Rate with decay | 1 | 0 | 1 |
| Response: Weighted rate | 0 | 0 | 0 |
| Dimension: No update | 7 | 0 | 0 |
| Dimension: S0 with decay | 0 | 0 | 3 |
| Dimension: Binary rate | 0 | 1 | 7 |
| Dimension: Rate with decay | 2 | 11 | 0 |
| Dimension: Weighted rate | 3 | 0 | 2 |

### S5. The role of the forgetting rule in the inter-trial effects

Our starting point updating rule, ‘S0 with full memory’, predicts that inter-trial effects would decrease in magnitude over the course of an experiment, as the precision of the prior increases and new evidence consequently has less of an effect. The ‘S0 with decay’ rule, on the other hand, does not make this prediction, and our model comparison results clearly show that this rule explains the data better. In this section, we tested whether the size of inter-trial effects do, in fact, decrease over time, in order to confirm whether this difference in predictions is the reason why the ‘S0 with decay’ rule performed better. We did this by splitting the sequence of trials in half for each participant and experiment and calculating inter-trial effects for repetition versus switch of the response-defining feature separately for the early and late half of each experiment (see Figure S4). In each experiment inter-trial effects were, in fact, slightly larger in the second half, although these differences were not statistically significant (Exp. 1: F(1,11)=2.28, p=0.16, BF=0.46; Exp. 2: F(1,11)=3.00, p=0.11, BF=0.50; Exp. 3: F(1,11)=0.45, p=0.52, BF=0.39). This contradicts the predictions of the ‘S0 with full memory’ rule and explains why the ‘S0 with decay’ rule performed better.

**Figure S4.**
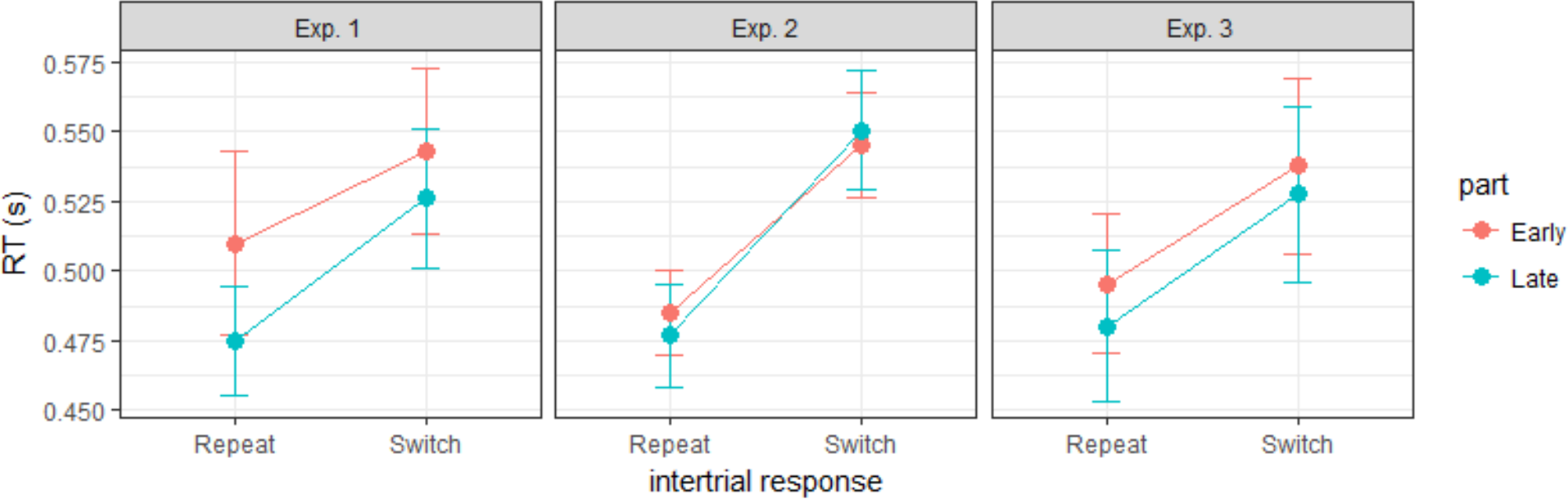
Effects of response feature repetition/switch on mean RTs in the early and late half of each experiment. Error bars show the standard error of the mean.

### S6. Comparison of drift rate updating rules in inter-trial effects

As pointed out in the Discussion section, our two different drift rate updating rules with a memory of more than a single trial back – ‘rate with decay’ and ‘weighted rate’ – are in general quite similar, but make opposite predictions regarding two types of sequences: a sequence of repeats followed by a switch (i.e., {R, …, R, S}) and an equally long sequence of only switches ({S,… S,S}).

The ‘rate with decay’ rule predicts a higher rate after the {R,…, R, S} sequence compared to the {S,…, S, S} sequence, because, according to this rule, repeating a dimension always increases the rate on future trials regardless of dimension. The ‘weighted rate’ rule, by contrast, predicts that the rate should be particularly low after the {R,…, R, S} sequence, because each repetition means more weight is assigned to the repeated dimension and consequently less weight to the other dimension which will be the target dimension after the switch at the end.

Because the ‘rate with decay’ for dimension-based updating best explained the data in Experiment 2, while the ‘weighted’ rate rule best explained the data in Experiment 3, we tested whether RTs after the kind of sequences of dimension repetition and switch differ in the predicted way between these experiments. In particular, we compared RTs after sequences of two dimension repeats and one switch (RRS) with RTs after a sequence of three switches (SSS).

**Figure S5.**
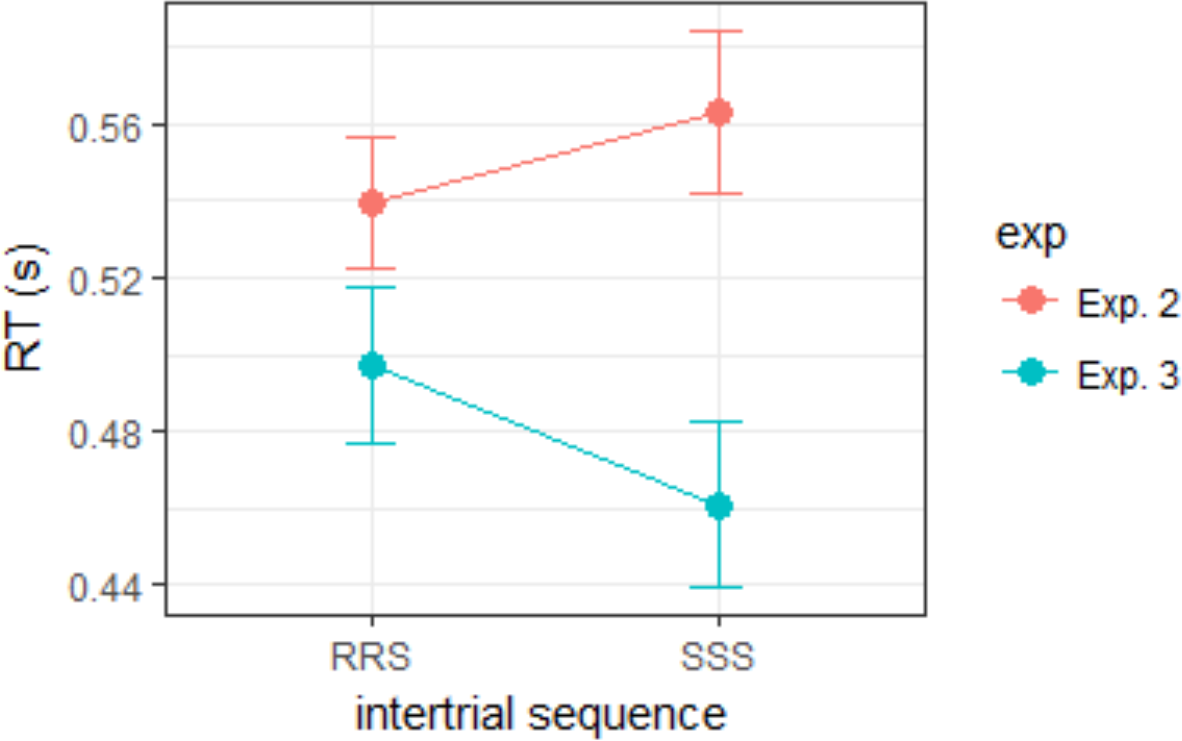
Mean RTs on switch trials at the end of different sequences of dimension repetitions (R) and switches (S) in Experiments 2 and 3. Error bars show the standard error of the mean.

Figure S5 shows the mean RTs at the end of these different sequences. In Experiment 2, RTs were significantly faster at the end of the RRS compared to the SSS sequence (t(11)=2.69, p<0.05), matching the prediction of the ‘rate with decay’ rule; in Experiment 3, by contrast, RTs were significantly faster at the end of the SSS compared to the RRS sequence (t(11)=2.3, p<0.05), following the prediction of the ‘(dimension-) weighted rate’ rule. Note that, in Experiment 2, the dimension was also the response-defining feature (RDF). Accordingly, the RTs for the different sequences of dimension repeats and switches could also have been influenced by starting-point updating based on the RDF. However, this is unlikely to explain the faster RTs at the end of RRS compared to SSS sequences, because the winning starting-point updating rule would make the opposite prediction. This implies that the effect of rate updating must have been greater than that of starting-point updating in Experiment 2, at least with regard to effects from more than one trial back (e.g., memory decay may have been slower for rate updates compared to starting-point updates). In addition, because the dimension was the RDF in Experiment 2, the rate updating based on dimension history in the winning model may partially reflect an effect of repetition/switch of the whole S-R link, rather than of just the dimension itself. This could explain the differential patterns seen in Experiment 2 vs. Experiment 3 (see main text for an elaboration of this argument).

### S7. Model fits to RT distributions

The LATER model in general fitted the RT distributions somewhat better than the DDM. In order to allow the reader to judge the quality of the fits for each model and the nature of the deviations, we here provide figures (Figure S6-S13) of individual subject RT distributions for each experiment, separately for color targets, orientation targets and target absent trials, and the LATER model and DDM fit to each distribution. We show these fits for models with no updating, because the effects of the updating rules only become relevant when taking trial history into account and do not improve the fit to the overall distributions, but with a non-decision time. The predictions of the DDM and the LATER model are in general quite similar, but the DDM tends to have a somewhat longer “tail”.

**Figure S6.**
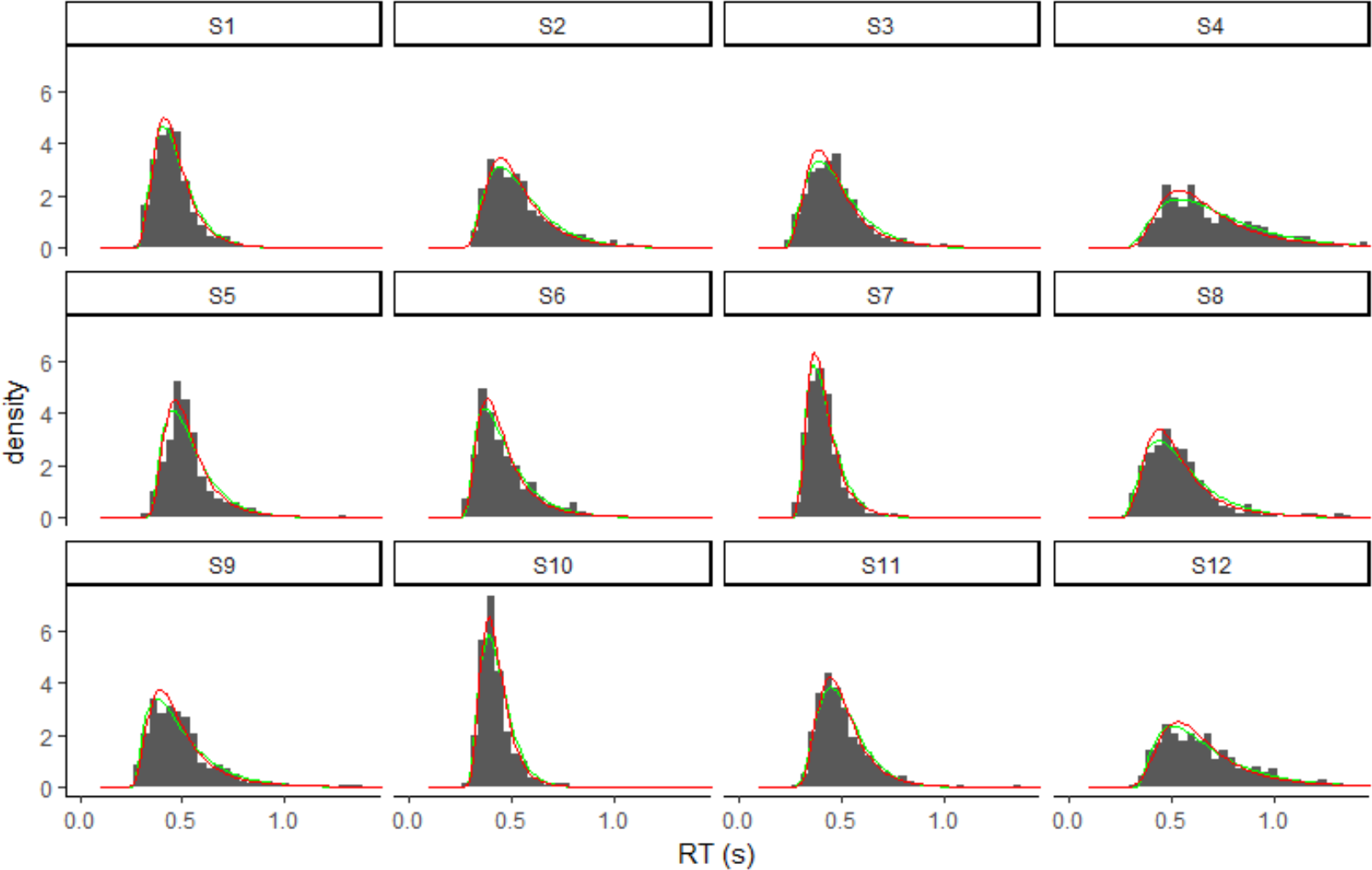
RT distributions for individual participants and model fits, without updating but with a non-decision time, for target absent trials in Experiment 1. The red line shows the fit of the LATER model while the green line shows the fit of the DDM.

**Figure S7.**
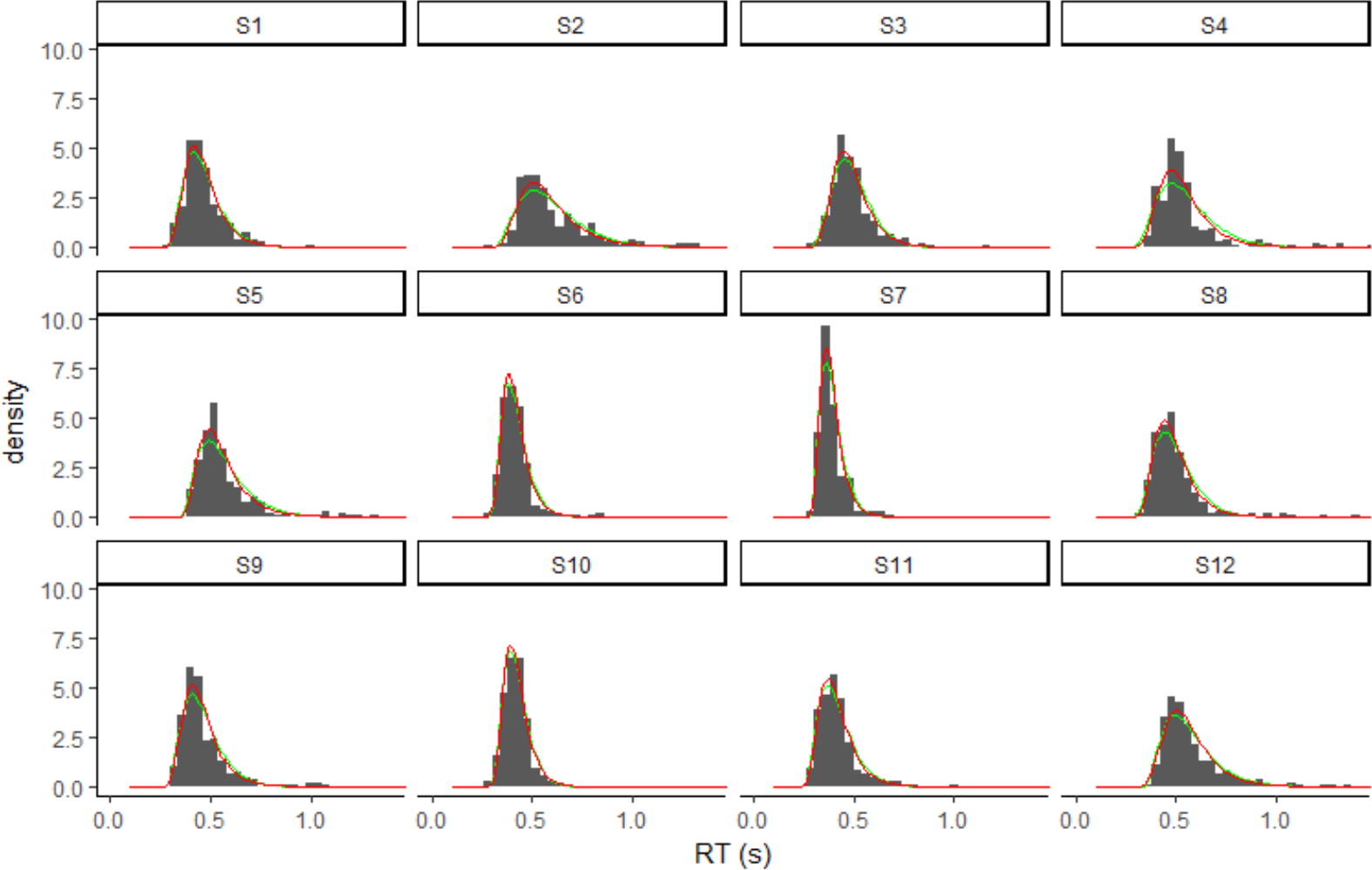
RT distributions for individual participants and model fits, without updating but with a non-decision time, for orientation target trials in Experiment 1. The red line shows the fit of the LATER model while the green line shows the fit of the DDM.

**Figure S8.**
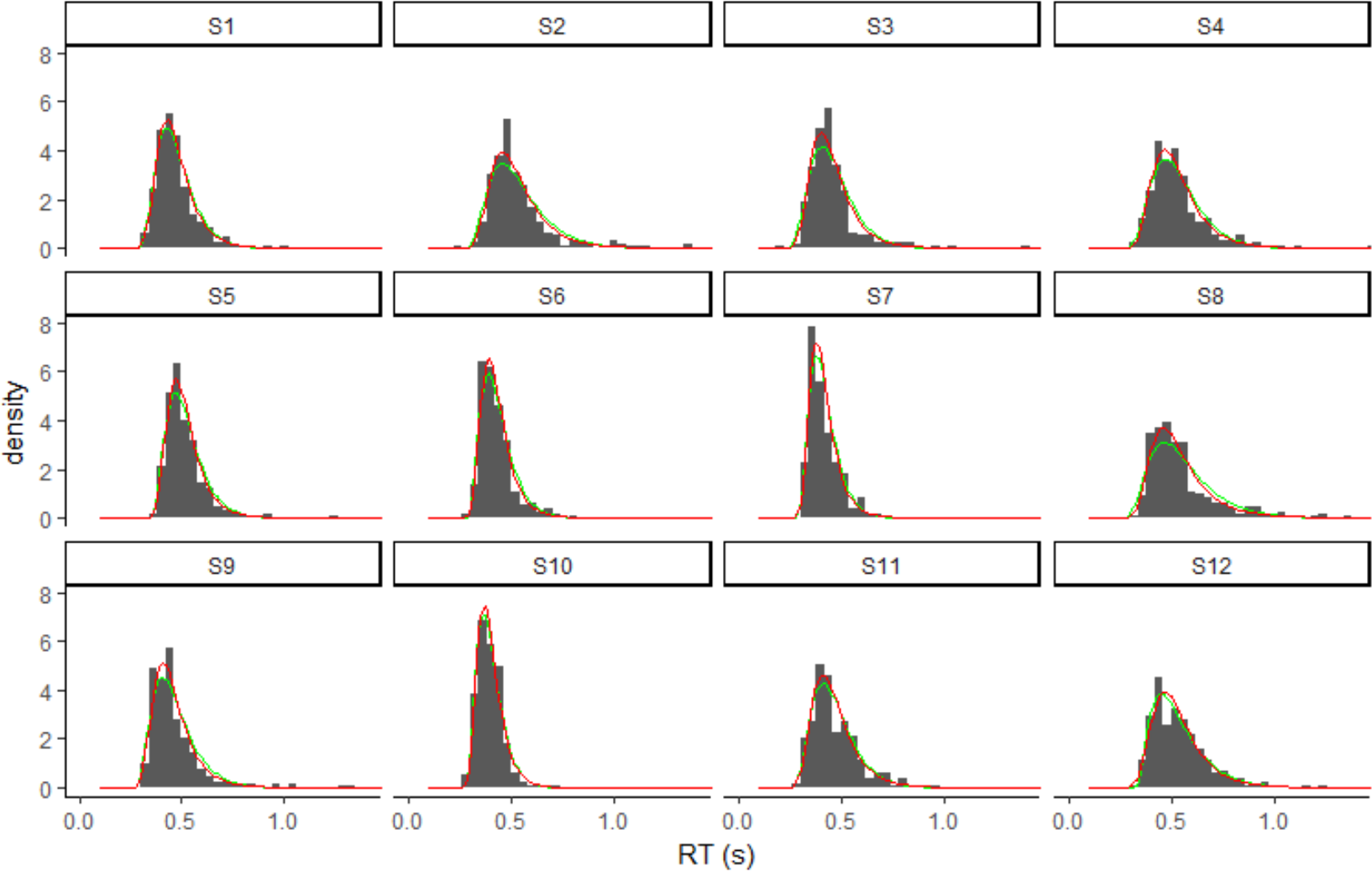
RT distributions for individual participants and model fits, without updating but with a non-decision time, for color target trials in Experiment 1. The red line shows the fit of the LATER model while the green line shows the fit of the DDM.

**Figure S9.**
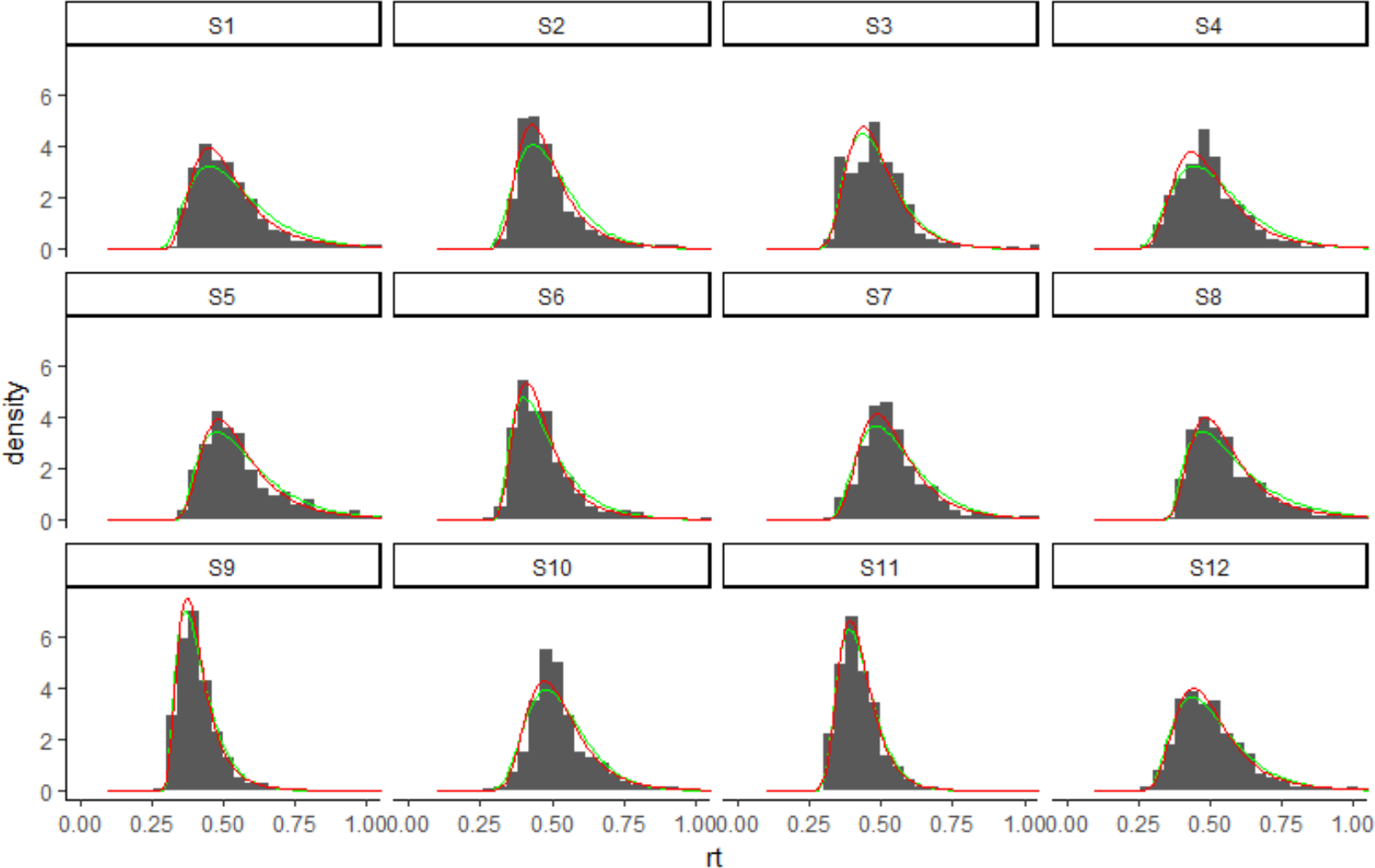
RT distributions for individual participants and model fits, without updating but with a non-decision time, for color target trials in Experiment 2. The red line shows the fit of the LATER model while the green line shows the fit of the DDM.

**Figure S10.**
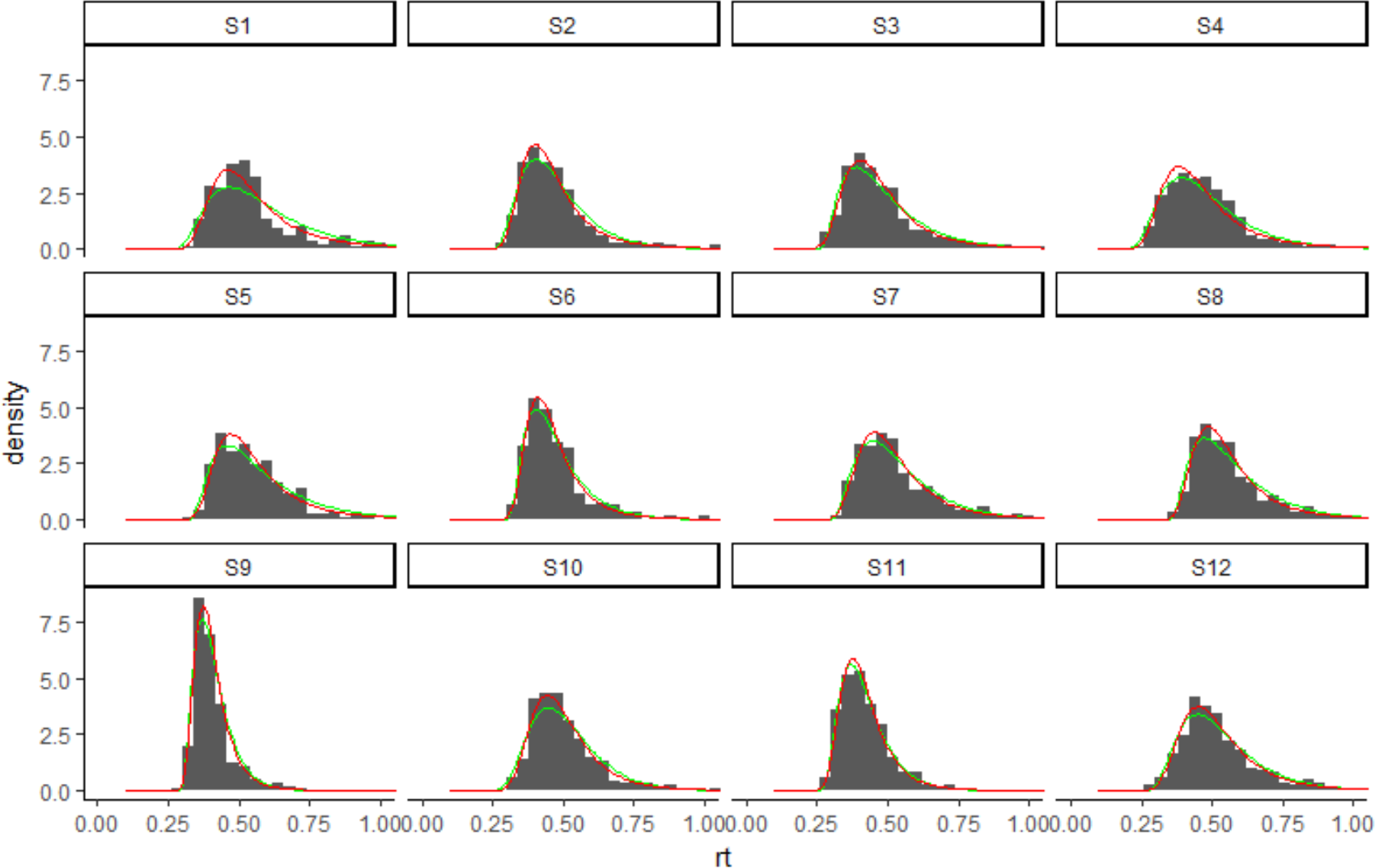
RT distributions for individual participants and model fits, without updating but with a non-decision time, for orientation target trials in Experiment 2. The red line shows the fit of the LATER model while the green line shows the fit of the DDM.

**Figure S11.**
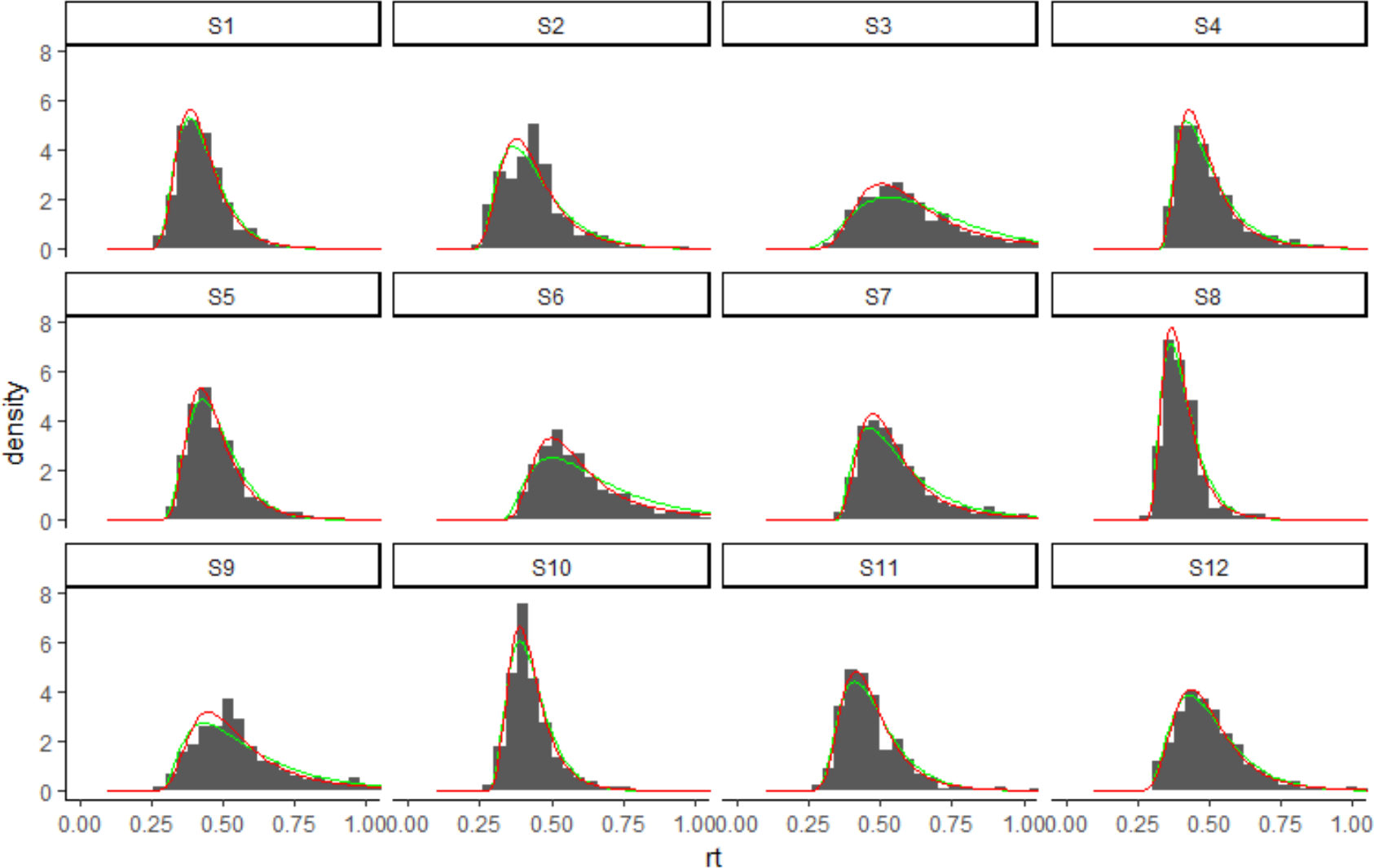
RT distributions for individual participants and model fits, without updating but with a non-decision time, for target absent trials in Experiment 3. The red line shows the fit of the LATER model while the green line shows the fit of the DDM.

**Figure S12.**
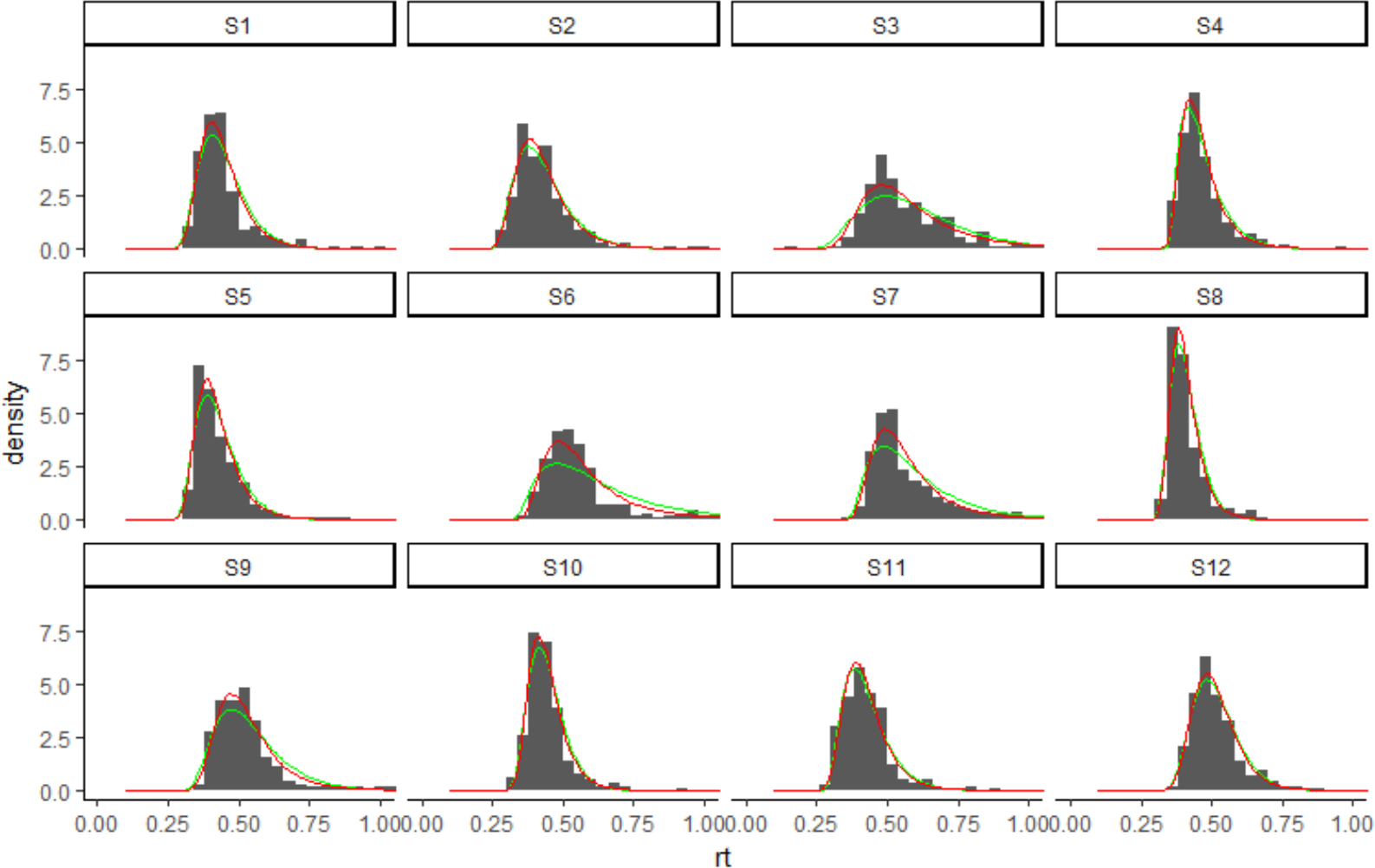
RT distributions for individual participants and model fits, without updating but with a non-decision time, for orientation target trials in Experiment 3. The red line shows the fit of the LATER model while the green line shows the fit of the DDM.

**Figure S13.**
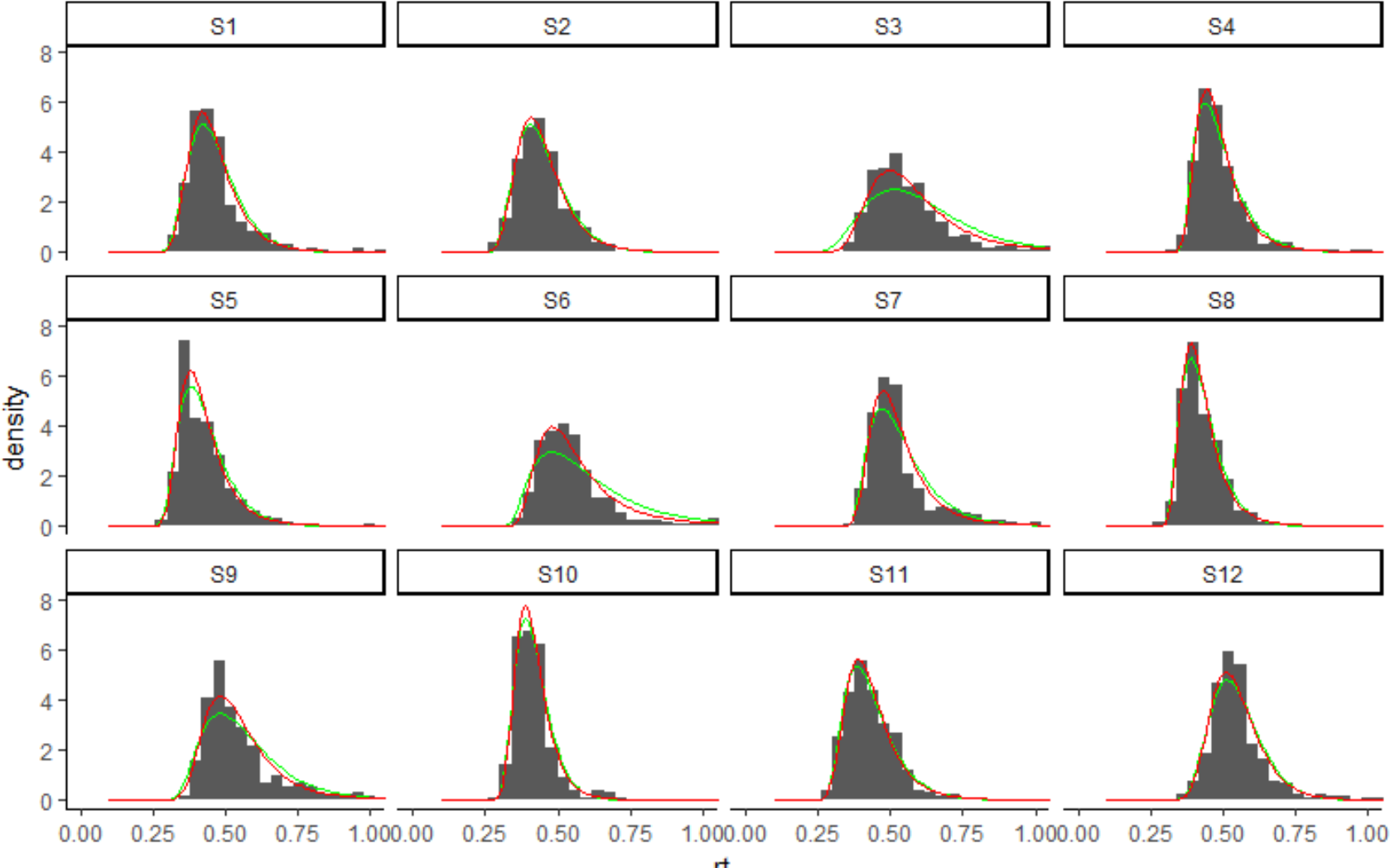
RT distributions for individual participants and model fits, without updating but with a non-decision time, for color target trials in Experiment 3. The red line shows the fit of the LATER model while the green line shows the fit of the DDM.

## Acknowledgements

This work was supported by German research foundation DFG MU773/16-1 to HJM and DFG SH166/3-1 to ZS. The authors would like to thank Lena Schröder and Kerstin Wenzel for recruiting and testing participants, two anonymous reviewers for their insightful comments on an earlier version of this paper, and Uta Noppeney for helpful discussions.

## References

1. Kristjánsson Á, Wang D, Nakayama K. The role of priming in conjunctive visual search. Cognition. 2002;85: 37–52. doi:10.1016/S0010-0277(02)00074-4

2. Bravo MJ, Nakayama K. The role of attention in different visual-search tasks. Percept Psychophys. 1992;51: 465–472. doi:10.3758/BF03211642

3. Maljkovic V, Nakayama K. Priming of pop-out: I. Role of features. Mem Cognit. 1994;22: 657–672. doi:10.3758/BF03209251

4. Huang L, Holcombe AO, Pashler H. Repetition priming in visual search: Episodic retrieval, not feature priming. Mem Cognit. 2004;32: 12–20. doi:10.3758/BF03195816

5. Maljkovic V, Nakayama K. Priming of pop-out: II. The role of position. Percept Psychophys. 1996;58: 977–991.

6. Found A, Müller HJ. Searching for unknown feature targets on more than one dimension: Investigating a “dimension-weighting” account. Percept Psychophys. 1996;58: 88–101. doi:10.3758/BF03205479

7. Müller HJ, Reimann B, Krummenacher J. Visual search for singleton feature targets across dimensions: Stimulus- and expectancy-driven effects in dimensional weighting. J Exp Psychol Hum Percept Perform. 2003;29: 1021–1035. doi:10.1037/0096-1523.29.5.1021

8. Müller HJ, Heller D, Ziegler J. Visual search for singleton feature targets within and across feature dimensions. Percept Psychophys. 1995;57: 1–17. doi:10.3758/BF03211845

9. Müller H, Krummenacher J, Heller D. Dimension-specific intertrial facilitation in visual search for pop-out targets: Evidence for a top-down modulable visual short-term memory effect. Vis Cogn. 2004;11: 577–602. doi:10.1080/13506280344000419

10. Müller HJ, Krummenacher J. Locus of dimension weighting: Preattentive or postselective? Vis Cogn. 2006;14: 490–513. doi:10.1080/13506280500194154

11. Lamy DF, Kristjánsson Á. Is goal-directed attentional guidance just intertrial priming? A review. J Vis. 2013;13: 14–14. doi:10.1167/13.3.14

12. Mortier K, Theeuwes J, Starreveld P. Response Selection Modulates Visual Search Within and Across Dimensions. J Exp Psychol Hum Percept Perform. 2005;31: 542–557. doi:10.1037/0096-1523.31.3.542

13. Rangelov D, Müller HJ, Zehetleitner M. Independent dimension-weighting mechanisms for visual selection and stimulus identification. J Exp Psychol Hum Percept Perform. 2011;37: 1369–1382. doi:10.1037/a0024265

14. Neill WT. Episodic Retrieval in Negative Priming and Repetition Priming. J Exp Psychol Learn Mem Cogn. 1997;23: 1291–1305.

15. Lamy D, Yashar A, Ruderman L. A dual-stage account of inter-trial priming effects. Vision Res. 2010;50: 1396–1401. doi:10.1016/j.visres.2010.01.008

16. Moran R, Zehetleitner M, Müller HJ, Usher M. Competitive guided search: Meeting the challenge of benchmark RT distributions. J Vis. 2013;13: 24–24. doi:10.1167/13.8.24

17. Wolfe JM, Palmer EM, Horowitz TS. Reaction time distributions constrain models of visual search. Vision Res. 2010;50: 1304–1311. doi:10.1016/j.visres.2009.11.002

18. Gold JI, Shadlen MN. Neural computations that underlie decisions about sensory stimuli. Trends Cogn Sci. 2001;5: 10–16. doi:10.1016/S1364-6613(00)01567-9

19. Gold JI, Shadlen MN. The Neural Basis of Decision Making. Annu Rev Neurosci. 2007;30: 535–574. doi:10.1146/annurev.neuro.29.051605.113038

20. Ratcliff R. Group reaction time distributions and an analysis of distribution statistics. Psychol Bull. 1979;86: 446–461. doi:10.1037/0033-2909.86.3.446

21. Ratcliff R, Smith P. A comparison of sequential sampling models for two-choice reaction time. Psychol Rev. 2004;111: 333–367. doi:10.1037/0033-295x.111.2.333

22. Smith PL, Ratcliff R. Psychology and neurobiology of simple decisions. Trends Neurosci. 2004;27: 161–168. doi:10.1016/j.tins.2004.01.006

23. Cohen A, Shoup R. Perceptual Dimensional Constraints in Response Selection Processes. Cognit Psychol. 1997;32: 128–181. doi:10.1006/cogp.1997.0648

24. Otto TU, Mamassian P. Noise and Correlations in Parallel Perceptual Decision Making. Curr Biol. 2012;22: 1391–1396. doi:10.1016/j.cub.2012.05.031

25. Carpenter RHS, Williams MLL. Neural computation of log likelihood in control of saccadic eye movements. Nature. 1995;377: 59–62. doi:10.1038/377059a0

26. Noorani I, Carpenter RHS. The LATER model of reaction time and decision. Neurosci Biobehav Rev. 2016;64: 229–251. doi:10.1016/j.neubiorev.2016.02.018

27. Hanes DP, Schall JD. Neural control of voluntary movement initiation. Science. 1996;274: 427–430.

28. Bitzer S, Park H, Blankenburg F, Kiebel SJ. Perceptual decision making: Drift- diffusion model is equivalent to a Bayesian model. Front Hum Neurosci. 2014;8. doi:10.3389/fnhum.2014.00102

29. Yu AJ, Cohen JD. Sequential effects: Superstition or rational behavior? Adv Neural Inf Process Syst. 2008;21: 1873–1880.

30. Anderson AJ, Carpenter RHS. Changes in expectation consequent on experience, modeled by a simple, forgetful neural circuit. J Vis. 2006;6: 5–5. doi:10.1167/6.8.5

31. van den Berg R, Awh E, Ma WJ. Factorial Comparison of Working Memory Models. Psychol Rev. 2014;121: 124–149. doi:10.1037/a0035234

32. Martini P. System identification in Priming of Pop-Out. Vision Res. 2010;50: 2110–2115. doi:10.1016/j.visres.2010.07.024

33. Kruijne W, Brascamp JW, Kristjánsson Á, Meeter M. Can a single short-term mechanism account for priming of pop-out? Vision Res. 2015;115: 17–22. doi:10.1016/j.visres.2015.03.011

34. Brascamp JW, Pels E, Kristjánsson Á. Priming of pop-out on multiple time scales during visual search. Vision Res. 2011;51: 1972–1978. doi:10.1016/j.visres.2011.07.007

35. Rangelov D, Müller HJ, Zehetleitner M. Failure to pop out: Feature singletons do not capture attention under low signal-to-noise ratio conditions. J Exp Psychol Gen. 2017;146: 651–671. doi:10.1037/xge0000284

36. Becker SI. The mechanism of priming: Episodic retrieval or priming of pop-out? Acta Psychol (Amst). 2008;127: 324–339. doi:10.1016/j.actpsy.2007.07.005

37. Caddigan E, Lleras A. Saccadic repulsion in pop-out search: How a target’s dodgy history can push the eyes away from it. J Vis. 2010;10: 9–9. doi:10.1167/10.14.9

38. Rangelov D, Müller HJ, Zehetleitner M. Visual search for feature singletons: Multiple mechanisms produce sequence effects in visual search. J Vis. 2013;13: 22–22. doi:10.1167/13.3.22

39. Liesefeld HR, Moran R, Usher M, Müller HJ, Zehetleitner M. Search efficiency as a function of target saliency: The transition from inefficient to efficient search and beyond. J Exp Psychol Hum Percept Perform. 2016;42: 821–836. doi:10.1037/xhp0000156

40. Krummenacher J, Müller HJ, Heller D. Visual search for dimensionally redundant pop-out targets: Redundancy gains in compound tasks. Vis Cogn. 2002;9: 801–837. doi:10.1080/13506280143000269

41. Töllner T, Gramann K, Müller HJ, Kiss M, Eimer M. Electrophysiological markers of visual dimension changes and response changes. J Exp Psychol Hum Percept Perform. 2008;34: 531–542. doi:10.1037/0096-1523.34.3.531

42. Töllner T, Rangelov D, Müller HJ. How the speed of motor-response decisions, but not focal-attentional selection, differs as a function of task set and target prevalence. Proc Natl Acad Sci. 2012;109: E1990–E1999. doi:10.1073/pnas.1206382109

43. Yang T, Shadlen MN. Probabilistic reasoning by neurons. Nature. 2007;447: 1075–1080. doi:10.1038/nature05852

44. Duncan J. Visual search and visual attention. Routledge; 1985. pp. 86–106.

45. Neill WT, Valdes LA, Terry KM, Gorfein DS. Persistence of negative priming: II. Evidence for episodic trace retrieval. J Exp Psychol Learn Mem Cogn. 1992;18: 993–1000.

46. Töllner T, Gramann K, Müller HJ, Eimer M. The anterior N1 component as an index of modality shifting. J Cogn Neurosci. 2009;21: 1653–1669. doi:10.1162/jocn.2009.21108

47. Töllner T, Müller HJ, Zehetleitner M. Top-Down Dimensional Weight Set Determines the Capture of Visual Attention: Evidence from the PCN Component. Cereb Cortex. 2012;22: 1554–1563. doi:10.1093/cercor/bhr231

48. Lee MD, Fuss IG, Navarro DJ. A Bayesian Approach to Diffusion Models of Decisionmaking and Response Time. Proceedings of the 19th International Conference on Neural Information Processing Systems. Cambridge, MA, USA: MIT Press; 2006. pp. 809–816. Available: http://dl.acm.org/citation.cfm?id=2976456.2976558

49. Itti L, Koch C. Computational modelling of visual attention. Nat Rev Neurosci. 2001;2: 194. doi:10.1038/35058500

50. Wolfe JM. When do I Quit? The Search Termination Problem in Visual Search. In: Dodd MD, Flowers JH, editors. The Influence of Attention, Learning, and Motivation on Visual Search. Springer New York; 2012. pp. 183–208. doi:10.1007/978-1-4614-4794-8_8

51. Chun MM, Wolfe JM. Just Say No: How Are Visual Searches Terminated When There Is No Target Present? Cognit Psychol. 1996;30: 39–78. doi:10.1006/cogp.1996.0002

52. Mozer MC, Shettel M, Vecera S. Top-Down Control of Visual Attention: A Rational Account. In: Weiss Y, Schölkopf PB, Platt JC, editors. Advances in Neural Information Processing Systems 18. MIT Press; 2006. pp. 923–930. Available: http://papers.nips.cc/paper/2875-top-down-control-of-visual-attention-a-rational-account.pdf

53. Wolfe JM, Butcher SJ, Lee C, Hyle M. Changing your mind: On the contributions of top-down and bottom-up guidance in visual search for feature singletons. J Exp Psychol Hum Percept Perform. 2003;29: 483–502. doi:10.1037/0096-1523.29.2.483

54. Bundesen C. A theory of visual attention. Psychol Rev. 1990;97: 523–547. doi:10.1037/0033-295X.97.4.523

55. Ásgeirsson ÁG, Kristjánsson Á, Bundesen C. Independent priming of location and color in identification of briefly presented letters. Atten Percept Psychophys. 2014;76: 40–48. doi:10.3758/s13414-013-0546-6

56. Ásgeirsson ÁG, Kristjánsson Á, Bundesen C. Repetition priming in selective attention: A TVA analysis. Acta Psychol (Amst). 2015;160: 35–42. doi:10.1016/j.actpsy.2015.06.008

57. Krummenacher J, Müller HJ, Zehetleitner M, Geyer T. Dimension- and space-based intertrial effects in visual pop-out search: modulation by task demands for focal-attentional processing. Psychol Res. 2008;73: 186–197. doi:10.1007/s00426-008-0206-y

58. Neill WT, Valdes LA. Persistence of Negative Priming: Steady State or Decay? J Exp Psychol Learn Mem Cogn. 1992;18: 565–576.

59. Brainard DH. The Psychophysics Toolbox. Spat Vis. 1997;10: 433–436. doi:10.1163/156856897X00357

60. Kleiner M, Brainard D, Pelli D. What’s new in Psychtoolbox-3? Perception. 2007;36: ECVP Abstract Supplement.

61. Brimijoin WO, O’Neill WE. Patterned Tone Sequences Reveal Non-Linear Interactions in Auditory Spectrotemporal Receptive Fields in the Inferior Colliculus. Hear Res. 2010;267: 96–110. doi:10.1016/j.heares.2010.04.005

62. de Bruijn NG. A Combinatorial Problem. Proc Akad Van Westeschappen. 1946;49: 758–764.

